# Adaptation of human iPSC-derived cardiomyocytes to tyrosine kinase inhibitors reduces acute cardiotoxicity via metabolic reprogramming

**DOI:** 10.1101/365841

**Authors:** Huan Wang, Robert P. Sheehan, Adam C. Palmer, Robert A. Everley, Sarah A. Boswell, Noga Ron-Harel, Alison E. Ringel, Kristina M. Holton, Connor A. Jacobson, Alison R. Erickson, Laura Maliszewski, Marcia C. Haigis, Peter K. Sorger

**Author notes:** Huan Wang, Institute of Systems Biomedicine, School of Basic Medical Sciences, Peking University Health Science Center, Beijing 10091, China.

## Abstract

Tyrosine kinase inhibitors (TKIs) are widely used to treat solid tumors but can be cardiotoxic. The molecular basis for this toxicity and its relationship to therapeutic mechanisms remain unclear; we therefore undertook a systems-level analysis of human cardiomyocytes exposed to four TKIs. Cardiomyocytes (CMs) differentiated from human induced pluripotent stem cells (hiPSCs) were exposed to sunitinib, sorafenib, lapatinib or erlotinib and responses assessed by functional assays, microscopy, RNA sequencing and mass spectrometry (GEO GSE114686; PRIDE PXD012043). TKIs have diverse effects on hiPSC-CMs distinct from inhibition of tyrosine-kinase mediated signal transduction; cardiac metabolism is particularly sensitive. Following Sorafenib treatment, oxidative phosphorylation is down-regulated, resulting in a profound defect in mitochondrial energetics. Cells adapt by upregulating aerobic glycolysis. Adaptation makes cells less acutely sensitive to Sorafenib, but may have long-term negative consequences. Thus, cardiomyocytes exhibit adaptive responses to anti-cancer drugs conceptually similar to those previously shown in tumors to mediate drug resistance.

## INTRODUCTION

As therapeutic responses to “targeted” anti-cancer drugs become increasingly sustained, drug-induced cardiotoxicity is a growing concern. Cardiotoxicity is observed with a wide range of drugs, including tyrosine kinase inhibitors (Chu et al., 2007), immune checkpoint inhibitors (Johnson et al., 2016; Moslehi et al., 2018), nonsteroidal anti-inflammatory drugs (Bresalier et al., 2005) and proteasome inhibitors (Waxman et al., 2018). In patients, exposure to these drugs causes one or more of the following adverse effects: hypertension, arrhythmia, decreased left ventricular ejection fraction (LVEF), myocarditis, cardiac ischemia and cardiac failure (Magdy et al., 2018). A dramatic example of cardiotoxicity is observed in pediatric patients treated with anthracycline chemotherapy plus radiation. Such patients have a 7-fold higher risk of death due to cardiovascular damage than age-matched controls (Mertens et al., 2008). Cardio-protective measures, such as drug holidays, dose reduction, and/or administration of β-adrenergic receptor or angiotensin-converting-enzyme (ACE) inhibitors are indicated following a greater than 10% decline in LVEF (or if the absolute value is < 53% (Gavila et al., 2017)). However, treatment for drug-induced cardiotoxicity is limited: beta blockers and ACE inhibitors relieve the physiological symptoms of adverse cardiac events but do not alter the molecular processes responsible for cardiotoxicity or the resulting tissue damage.

Anthracycline-mediated cardiotoxicity and drug-induced arrhythmias caused by inhibition of the hERG potassium channel (product of the KCNH2 gene) are two well studied forms of cardiotoxicity. The anthracycline doxorubicin induces double-stranded breaks in DNA (Zhang et al., 2012) and cumulative drug dose is a strong predictor of risk for congestive heart failure (Von Hoff et al., 1979). The hERG potassium channel controls cardiac repolarization (Roden, 2004; Vandenberg et al., 2012) and hERG inhibition causes long QT syndrome and potentially *torsade de pointes*, an acutely life-threatening form of ventricular tachycardia; hERG is therefore an important anti-target for drug discovery.

Tyrosine kinase inhibitors (TKIs) are prototypical targeted anti-cancer therapies associated with varying degrees of cardiotoxicity different from anthracyclines and hERG-blockers. Both selective and non-selective TKIs are associated with cardiotoxicity, but selectivity is not obviously related to the magnitude of adverse effect (Orphanos et al., 2009). Among the drugs studied in this paper, Sunitinib (Suntent^®^) is most often associated with clinically detectable cardiotoxicity, including hypertension (8-47% of patients undergoing clinical trials), symptomatic depression of LVEF (9-28% in trials), and congestive heart failure (up to 8% in trials) (Chu et al., 2007). Sorafenib (Nexavar^®^) may cause hypertension (8-40% in trials) and less frequently cardiac ischemia (2-3% in trials) (Bhargava, 2009). Cardiomyopathies induced by Sunitinib and Sorafenib arise after prolonged time on drug (typically months) and are reversible following drug discontinuation (Francis et al., 2010; Schmidinger et al., 2008; Uraizee et al., 2011). In contrast, Lapatinib (Tykerb^®^)-associated cardiotoxicity is sporadic, less severe when it occurs, and most often associated with asymptomatic decreases in LVEF (Geyer et al., 2006). Erlotinib (Tarceva^®^) is very rarely associated with cardiac side-effects (Orphanos et al., 2009) and was included in our study as a bioactive, cardiotoxicity-negative control.

Previous *in vitro* and *in vivo* studies have established that TKIs can cause cardiotoxicity and hypertension through both on-target and off-target effects (Chen et al., 2008). Sunitinib and Sorafenib inhibit multiple receptor tyrosine kinases including VEGF Receptor (KDR) which is a known cause of hypertension (Schmidinger et al., 2008) albeit one that can usually be managed clinically (Robinson et al., 2010). Sunitinib has also been shown to induce cardiomyocyte apoptosis via inhibition of AMPK phosphorylation, an off-target effect (Kerkela et al., 2009). In addition, cardiomyocyte development and survival is dependent on many of the pathways inhibited by TKIs (e.g. PDGFRβ, VEGFR2, RAF1, etc.) although evidence that this is involved in toxicity remains limited (Force et al., 2007; Kerkela et al., 2006). The impact of duration of drug exposure on cardiotoxicity is not well understood. In cultured cells, the acute and long-term effects of anti-cancer drugs often differ, with the former reflecting direct target inhibition and the latter drug-induced senescence, death or adaptation. Adaptive responses often result in drug tolerance (Sun et al., 2014) but whether this occurs in cardiomyocytes is unknown.

In this study we profiled phenotypic and molecular responses to four widely used TKIs (Sunitinib, Sorafenib, Lapatinib and Erlotinib) in human induced pluripotent stem cell-derived cardiac myocytes (hiPSC-CMs) across varying doses of drug and times of exposure. hiPSC-CMs are an increasingly popular culture system in which to study cardiac biology because they are readily available, genetically well-defined, and more representative of human biology than immortalized cell lines or rodent models. However, hiPSC-CMs are less mature than fully-differentiated primary human cardiomyocytes and do not have the well-formed sarcomeres characteristic of adult cells (Burridge et al., 2016; Sharma et al., 2017). Systematic profiling of hiPSC-CMs was initiated with three goals: (i) to acquire matched proteomic and transcriptomic datasets of TKI exposure across a range of drug doses and exposure times to determine how toxicity relates to these variables, (ii) to investigate at least one prominent phenotype functionally and demonstrate the utility of the overall dataset (iii) to inform future counter-screening in drug discovery by establishing the reliability of different molecular profiling technologies across multiple batches of hiPSCs. For the four TKIs studied, RNA-Seq and proteomics of hiPSC-CMs revealed widespread changes in cardiac metabolism and disruption of sarcomere proteins – the magnitude of these changes correlated with clinical cardiotoxicity.

## RESULTS

### Phenotypic responses to four TKIs in human iPSC-derived cardiomyocytes

Cor.4U hiPSC-CMs were obtained from the manufacturer (Axiogenesis; now Ncardia). The cells underwent directed differentiated from iPSCs, generated from a human female, into cardiac myocytes. Cor.4U cells were plated in vendor-specified growth medium containing 10% FBS for three days, switched to minimal medium with 1% FBS and TKIs were then added; cells were fed periodically with fresh serum-containing medium and samples withdrawn for analysis at varying times (Fig. 1A). The differentiation of iPSC-CMs is known to be heterogeneous and cultures contain cells at different developmental stages (Karakikes et al., 2015). We assessed the maturity of Cor.4U hiPSC-CM cultures by immunofluorescence microscopy using sarcomeric proteins. Two days after plating, >90% of cells stained positive for α-actinin and the cardiac isoform Troponin T in regular and alternating sarcomere striations, a characteristic of functional cardiomyocytes (Fig. 1B and Supplementary Fig. 1). The remaining ~10% of cells appeared to cardiomyocyte progenitors. Both classes of cells exhibited the spontaneous contraction and calcium pulsing typical of cardiomyocytes in culture (Blinova et al., 2017).

**Figure 1:**
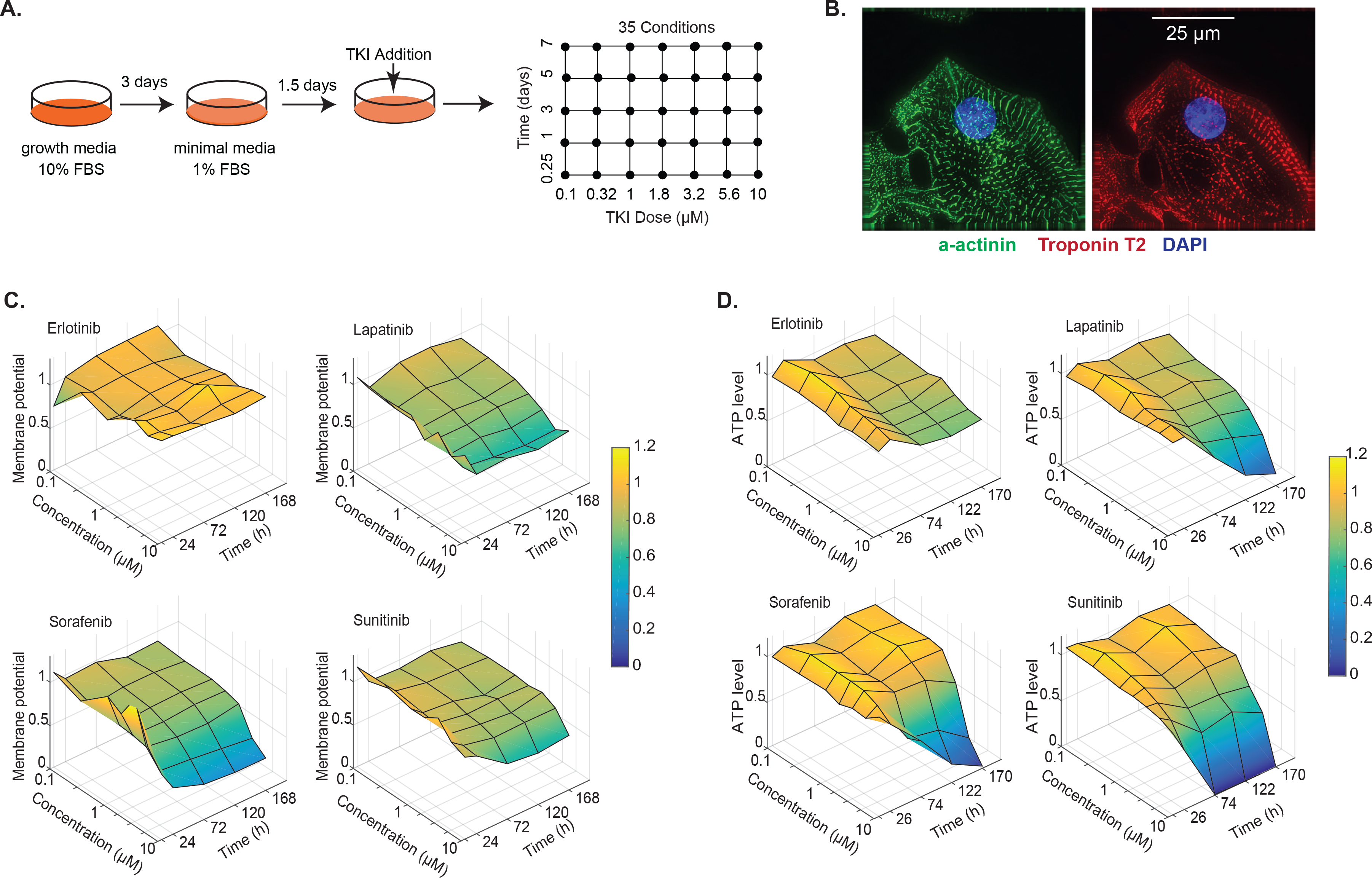
Phenotypic responses of human iPSC-derived cardiomyocytes to four TKIs. (**A**) Cor.4U hiPSC-CMs cells were cultured in vendor-specified growth media plus 10% FBS for 3 days, switched into a second vendor-specified media plus 1% FBS for 1.5 days and then exposed to drugs. For phenotypic studies, TKI doses between 0.1 −10 μM and treatment times between 6 hours and 7 days were examined (35 conditions in total). (**B**) hiPSC-CMs were stained with ⍺-actinin (green), Troponin T2 (red) and DAPI for nuclei (blue) and imaged by DeltaVision Elite. Images show a representative field from three independent experiments (scale bar is 25 μM; see Fig. S1 for lower magnification). (**C**) Mitochondrial membrane potentials of iPSC-CMs across 35 conditions for four TKIs. Values were normalized to cells treated with DMSO alone for the same amount of time. (**D**) ATP levels of whole-cell extracts measured using the CellTiter-Glo^®^ assay; conditions same as in Panel C.

In patients, the four TKIs used in this study elicited different degrees of cardiotoxicity on different time scales (Supplementary Table 1). The dose and time of TKI exposure *in vitro* that would generate data relevant to humans is not self-evident: cells in culture are sufficiently different from cells *in vivo* that the clinical maximum plasma concentration (C_max_) is only a rough guide to active concentrations of drug. C_max_ values for the four TKIs analyzed in this paper are reported to range from 0.03 to 18 μM with considerable variability (Supplementary Table 2), which likely arises from inter-individual differences in adsorption, distribution, metabolism, and excretion. We therefore took an empirical approach using two phenotypic assays. Changes in cellular ATP levels were measured as a function of drug dose and time using the CellTiter-Glo^®^ assay (which is often considered a proxy for cell viability) and mitochondrial membrane potential was measured using tetramethylrhodamine ethyl ester perchlorate (TMRE) followed by fluorescence imaging. hiPSC-CMs were exposed to each TKI at 1 of 8 drug doses ranging from 0.01 to 10 μM separated by half-log (~3.2 fold) or quarter-log (~1.8-fold) steps; the time of exposure ranged from 6 to 24, 72 or 168 hr (Fig. 1C). We found that Lapatinib, Sorafenib, and Sunitinib induced death in hiPSC-CMs at doses > 3 μM and exposure times over 5 days but that mitochondrial membrane potential was reduced at lower doses and shorter times (Fig. 1C). Erlotinib had little effect on mitochondrial membrane potential (Fig. 1C) or cell viability under any condition tested (Fig. 1D). Spontaneous beating of hiPSC-CMs was strongly inhibited by Sorafenib, Sunitinib and weakly by Lapatinib over similar concentration and time ranges as mitochondrial dysfunction was observed (Supplementary Fig. 1B). These data demonstrate the existence of a fairly wide exposure window in which TKIs induce a strong phenotype but cells remain viable and can therefore be subjected to molecular profiling. Moreover, the rank order of toxicity for the four TKIs on hiPSC-CMs was the same as previously reported for cardiotoxicity in human patients.

### Changes in RNA expression induced by exposure of hiPSC-CMs to TKIs

To identify gene expression changes associated with TKI exposure, we systematically varied dose and time centered on the active drug concentration identified by measurement of mitochondrial membrane potential and spontaneous beating. The resulting cruciform design of six conditions (plus one contemporaneous no-treatment control) covered dose and time with a minimum number of samples (Fig. 2A). Cells were exposed to one of four TKIs at 3 μM for 6, 24, 72 or 168 hr and also at 1 μM and 10 μM for 24 hr. Each condition was assayed in biological triplicate, yielding 18 samples per drug (84 overall including DMSO-treated controls; available in GEO as GSE114686). Performing a molecular profiling experiment at this scale required us to use multiple lots of cells, potentially introducing batch effects.

**Figure 2:**
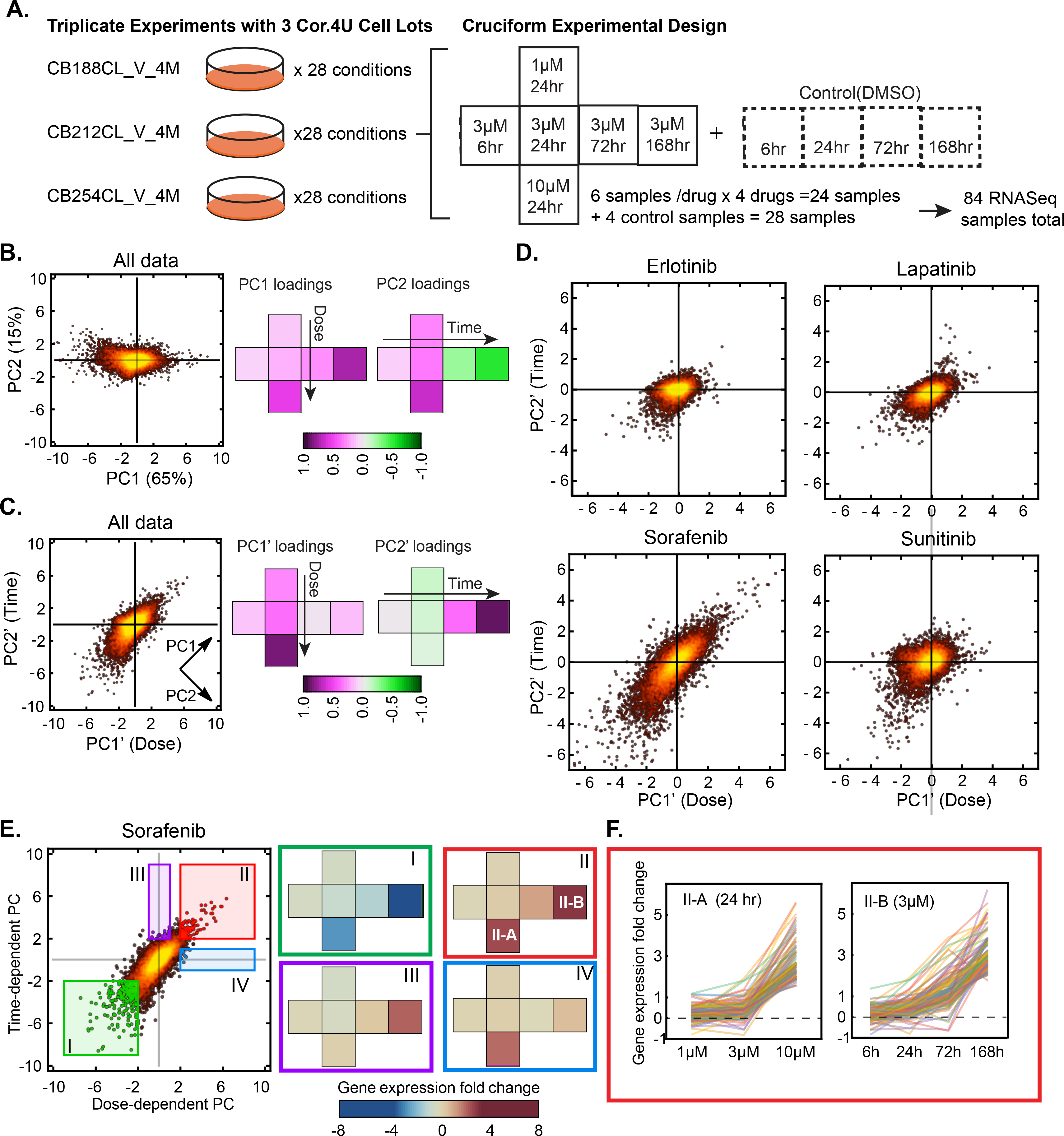
Responses of hiPSC-CMs to four TKIs as measured by RNA-Seq. (**A**) Three independent experiments with different lots of Cor.4U cells were performed. Drug treatment followed a cruciform design of 1, 3 or 10 μM for 24 hr and 3 μM drug for additional three time points (6, 72 and 168 hr) plus a no-treatment control denoted by a dashed box. 84 RNA samples were sequenced on a HiSeq 2500 yielding ~13,000 mapped transcripts per sample. (**B**) Principal component analysis (PCA) on fold-change values from RNA-Seq. Right: loadings of each experimental condition of the cruciform design. (**C**) The same data as in B with the axes rotated by ɵ=45° with was observed to optimally place dose along PC1’ and time along PC2’ (see loadings to right); inset shows the directions of the original PCs. (**D**) Scores of drug-induced gene expression changes were plotted against PC1’ and PC2’. (**E**) For the Sorafenib data in Panel D, genes corresponding to different dose-time regimes were selected (colored boxes I to IV) and the average gene expression changes were plotted on the experimental design. (**F**) Representative changes in expression for genes falling in boxes II-A and II-B from Panel E; each line represents one gene and colors are arbitrary. (See also Figure S2)

Sequencing data were normalized for time in culture by calculating expression fold changes relative to time-matched vehicle-only controls. In control cells, ~12,000 coding transcripts were detected; Pearson correlation coefficients for replicate assays of cells exposed to 10 μM Sorafenib or to vehicle-alone for 24 hr were 0.96-0.99, demonstrating good reproducibility across batches of cells (Supplementary Fig. 2). In the absence of TKI exposure, ~400-500 genes changed 1.5-fold or more in expression (FDR ≤ 0.05) over a period of seven days (Supplementary Fig. 2B). Genes corresponding to the GO term “cell cycle” were the most frequently down-regulated (Supplementary Fig. 2B). This is counter-intuitive since adult cardiomyocytes are terminally differentiated, non-mitotic cells. To investigate the cell cycle status of hiPSC-CMs in the presence and absence of TKIs we performed immunofluorescence imaging using the cardiomyocyte differentiation markers Mlc-2v and Mlc-2a (myosin light chains) and the S-phase marker phosphorylated retinoblastoma protein (p-pRb). We found that ~27% of Mlc-2v or Mlc-2a positive and ~17% of Mlc-2v or Mlc-2a negative cells were positive for p-pRb (Supplementary Fig. 2C), consistent with previous data showing that ~20% of hiPSC-CMs produced by directed differentiation express markers of cell cycle progression (Funakoshi et al., 2016; Shadrin et al., 2017; Zhou et al., 2017). The fraction of p-pRb^+^ cells did not change following 5 days of grown in the absence of drug, but was reduced ~22% after 3 days of treatment with Sorafenib at 3 μM (Supplementary Fig. 2C). Thus, downregulation of genes matching GO terms for “cell cycle” and related functions is likely a result of the known anti-mitogenic activity of TKIs. These changes are probably not relevant to adult cells in human patients and instead appear to reflect a limitation of hiPSC-CMs in culture as a model system.

When the gene expression fold-change data were combined and principal component analysis (PCA) performed, the first two principal components (PC1 and PC2) explained ~80% of variance; this represents excellent performance for PCA (Fig. 2B). By rotating the axes of the PCA plot (which preserves the data but can help with interpretation) we found that the new PC1’ corresponded primarily to differences in drug dose and PC2’ to differences in time of exposure (Fig. 2C). This is easily visualized by projecting PC1’ and PC2’ loadings onto the experimental design, as shown to the right of the PCA plots. When we generated rotated PCA plots for the four TKIs individually (Fig. 2D), we found that the correlation between dose and time was highest for Sorafenib. For example, in Figure 2E genes repressed by high drug doses at short times or by lower drug doses at longer times are found in the lower left quadrant (denoted by a green box) whereas genes induced at high drug doses and short times or lower drug doses and longer times were found in the upper right quadrant (red box). For Sorafenib, the magnitude of gene induction under the two sets of conditions was similar (Fig. 2F; data for Sunitinib are shown in Supplementary Fig. 2D).

To RNA-Seq data using an independent method, 90 differentially expressed genes selected from a range of GO categories were analyzed using quantitative real-time PCR (qRT-PCR; Supplementary Table 3). Three conditions (3 μM drug for 24 hr and 72 hr and 10 μM drug for 24 hr) were assayed in triplicate, for a total of 42 samples. We observed a Pearson correlation for differential gene expression of 0.93 to 0.98 depending on the drug (Supplementary Fig. 2E). Overall, these data show that patterns of TKI-induced gene expression are reproducible over separate lots of hiPSC-CMs and two different measurement methods. Moreover, for Sorafenib, and to a lesser extent for Sunitinib, it was the cumulative effect of dose and time that determined the magnitude of response, rather than dose or time individually. This observation increases the likelihood that analysis of cultured cells over short time periods (days) will be relevant to understanding longer duration exposure in patients (weeks to months).

### Biological processes altered by TKI exposure

To identify biological processes altered by TKI exposure we analyzed RNA-Seq data using G-means clustering followed by *goseq* enrichment analysis. G-means clustering (Hamerly and Elkan, 2004) is an abstraction of k-means clustering that detects the optimal number of clusters using a hierarchical approach: data in the neighborhood of a cluster are tested for fit to a Gaussian distribution; if the data have a more complex distribution, the cluster is successively split until a Gaussian is achieved. When RNA-Seq data were pooled, G-means clustering identified 16 major clusters, 13 of which were associated with significantly enriched GO terms. Two clusters shared the same GO term of “GO 0000278 Mitotic Cell Cycle” and were combined (Supplementary Table 4). Many other GO terms were associated with metabolism (*Amino acid metabolic process* GO:0006520, *Lipid biosynthetic process* GO:0008610, *Mitochondrion* GO:0005739) and some with contraction of muscle (*Sarcomere* GO:0030017). Some GO terms were enriched for a specific drug (e.g. GO: 0044424, GO:0042254, GO:0005739 by Sorafenib only) but others were enriched for all drugs (e.g. GO:0005615 down-regulated and GO:000861 upregulated). Within these categories we found multiple genes previously implicated in cardiac biology or dysfunction. For example, elevated steroid biosynthesis has been described in cases of cardiac hypertrophy with progression to heart failure (Ohtani et al., 2009).

Given that RNA-Seq data were acquired for multiple drugs, doses, and times of treatment, a fairly diverse and complex set of changes is to be expected. To begin to identify biological processes commonly altered by TKI treatment, we selected the top 1-2 GO terms in each cluster and plotted the log2-fold change for the top 20 genes in each GO term against the cruciform time-dose experimental design (Fig. 3). In the case of Sorafenib, expression changes observed at high dose and long times were consistent (recapitulating the data in Fig. 2), with *Sarcomere* GO:0030017 strongly down-regulated and *Ribosome biogenesis* GO:0042254 and *Cellular Amino Acid Metabolic Process* GO:0006520 strongly upregulated. When our data were combined with signatures previously published on exposure of cardiac cells to Doxorubicin (obtained from the NCBI Gene Expression Omnibus (GEO) database), PCA showed that the effects of TKIs and Doxorubicin were distinct, with TKI-driven responses and Doxorubicin-driven responses falling largely along different principle components (Supplementary Fig. 3). Thus, while TKIs differ in their effects on hiPSC-CMs, they are similar in causing differential regulation of genes associated with metabolism, biosynthetic processes and contractility; these are distinct from GO terms such as DNA damage associated with anthracycline exposure (Burridge et al., 2016).

**Figure 3:**
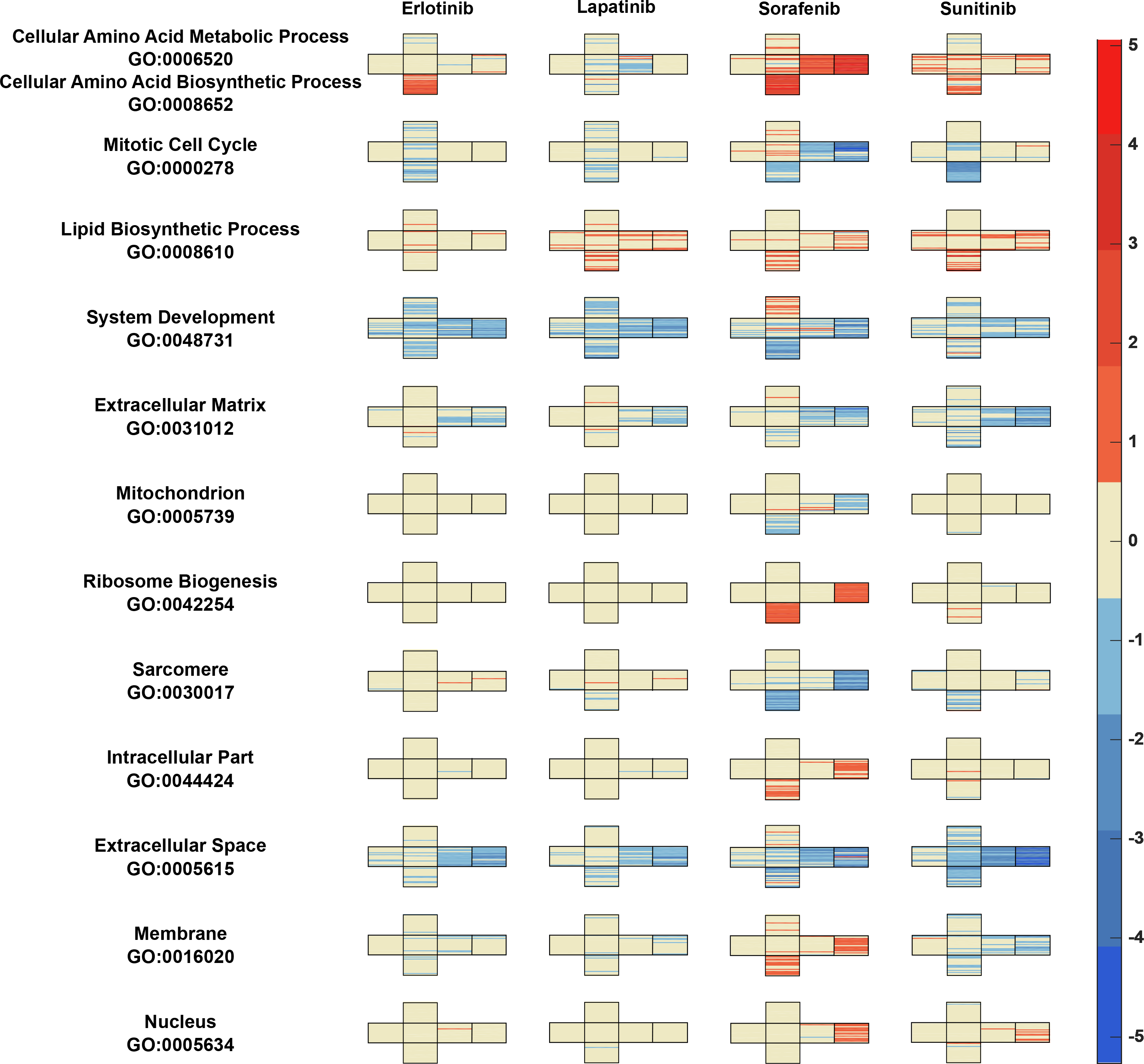
Effect of TKIs on the transcriptome across dose and time. Gene expression data were clustered by G-means clustering, which yielded 16 clusters, 13 of which had enriched GO terms (2 clusters with the same enriched GO term were combined into one). The top GO term(s) is shown for each cluster, with 20 representative genes from that cluster plotted in each box of the cruciform experimental design in Fig. 2A. Color scale is based on log2-transformed fold-changes. (See also Figure S3, Table S3)

### Changes in protein levels following exposure of hiPSC-CMs to TKIs

TKI-induced changes in the proteome were monitored by mass spectrometry following labelling with 10 different isobaric Tandem Mass Tags (TMTs) (Dayon and Sanchez, 2012). Cells were exposed to 3μM TKI for 24 or 72 hr and data from two batches of cells were run as three separate TMT experiments (30 samples total); these were combined by normalizing each drug treatment to a contemporaneous vehicle-only control (see Supplementary Fig. 4A and STAR METHOD). A total of ~7,700 proteins were detected in the union of the three experiments. PCA analysis showed that PC1 distinguished between early and late samples, whereas PC2 distinguished between Sorafenib, Sunitinib, and Lapatinib. Erlotinib induced relatively small deviations from the DMSO-only control and replicate samples for all drugs clustered close to each other, demonstrating good reproducibility (Fig. 4A; note however, individual protein levels in the proteomic data are subject to some batch effects, see Supplementary Fig. 4B). Differentially regulated proteins (relative to the vehicle-only negative control for the corresponding time point) were identified using a one-way ANOVA test corrected for multiple hypothesis testing (FDR ≤ 0.05 and fold change ≥ 1.2). In the case of Sunitinib, 562 differentially expressed proteins were detected at t = 72 hours in contrast to 90 proteins for Erlotinib-treated cells (Supplementary Fig. 4C). The data were clustered by K-means clustering with k = 11; adding more clusters did not significantly reduce the total within-cluster sum of squares (Supplementary Fig. 4D). We then analyzed the enriched GO terms for each cluster (Supplementary Table 5). Proteins in several clusters were drug-specific; for example, proteins in Cluster 1 (*Peroxisome* GO:0005777) were down-regulated by Lapatinib alone; proteins in Cluster 10 (*Sarcomere* GO:0030017) were down-regulated following exposure to Sorafenib or Sunitinib; and proteins in Cluster 11 (*Respiratory electron transport chain* GO:0022904) were up-regulated by Sorafenib and Sunitinib (Fig. 4B). Terms associated with mitochondrion, oxidation-reduction, alpha amino acid metabolism, sarcomere, DNA strand elongation, and cholesterol biosynthetic process, were differentially regulated in both RNAseq and proteomics data (Fig. 4C).

**Figure 4:**
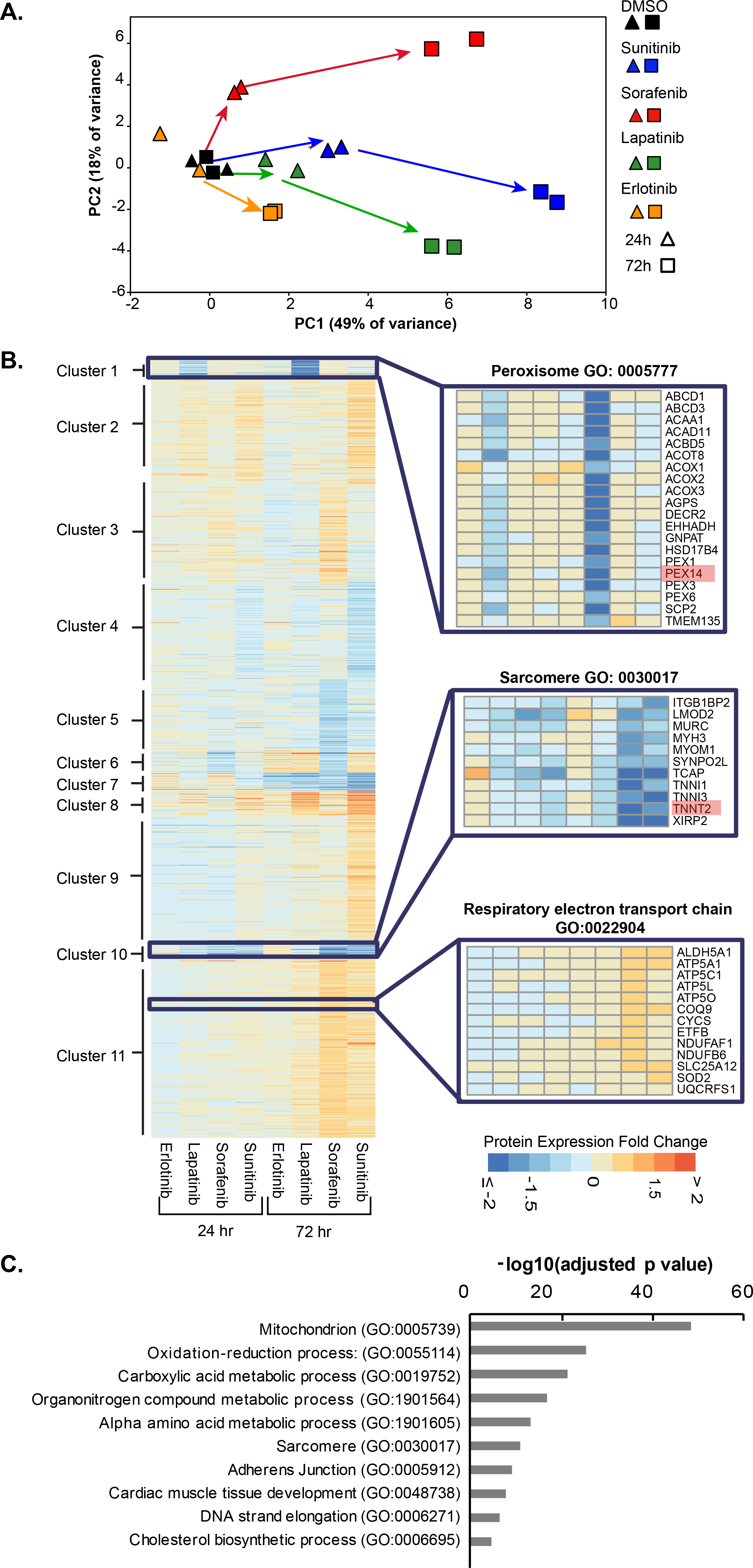
Effect of TKIs on the proteome across dose and time. **(A)** PCA of proteome changes for four TKIs at two time points; Data were pooled after calculating log2-transformed fold-changes in protein levels relative to DMSO at the corresponding time point and drugs were treated at 3 μM. Arrows depict the direction of change over time associated with each drug. (**B**) Proteomic fold-change data were clustered by k-means clustering with k=11; the top GO terms for each cluster were shown in Supplementary Table 6. Magnified views to right show individual genes for three enriched GO terms in these clusters. (**C**) GO terms enriched for genes that were significantly regulated at both RNA and protein levels by any drug; redundancy in the list of GO terms was eliminated to simplify the representation. (See also Figure S4, Table S5, Table S6)

### Integrating mRNA and proteomic data

TK Is typically inhibit multiple enzymes with similar affinity; Sorafenib and Sunitinib have particularly broad poly-pharmacology. Of the 537 human kinases in KinMap (Eid et al., 2017), ~75% (416) were detected in Cor.4U hiPSC-CMs: 274 kinases at the level of both mRNA and protein, 136 by mRNA alone and 6 by protein alone (Supplementary Table 6; whether a particular mRNA or protein is detected is a reflection of the sensitivity of the two assays, the ability of a specific peptide to fly in the mass spec, mRNA amplification efficiency etc., explaining any observed difference in the numbers of kinases detected by the two methods). Kinases were also among the genes or proteins differentially regulated by exposure to any one of the four TKIs – in some cases, the drug target was among the differentially expressed proteins. For example, the ERBB2 protein is a primary target of Lapatinib and its levels increased in cells treated with Lapatinib whereas the levels of VEGFR1 (FLT1) protein fell in response to Sorafenib treatment. By comparing our data to profiling studies that measure on- and off-target binding of TKIs to kinases (Davis et al., 2011), we estimate that ~143 potential targets for Sunitinib are expressed in Cor.4U hiPSC-CMs as are 25 potential targets for Sorafenib, 5 for Lapatinib and 20 for Erlotinib (Fig, 5A, Supplementary Table 7).

**Figure 5:**
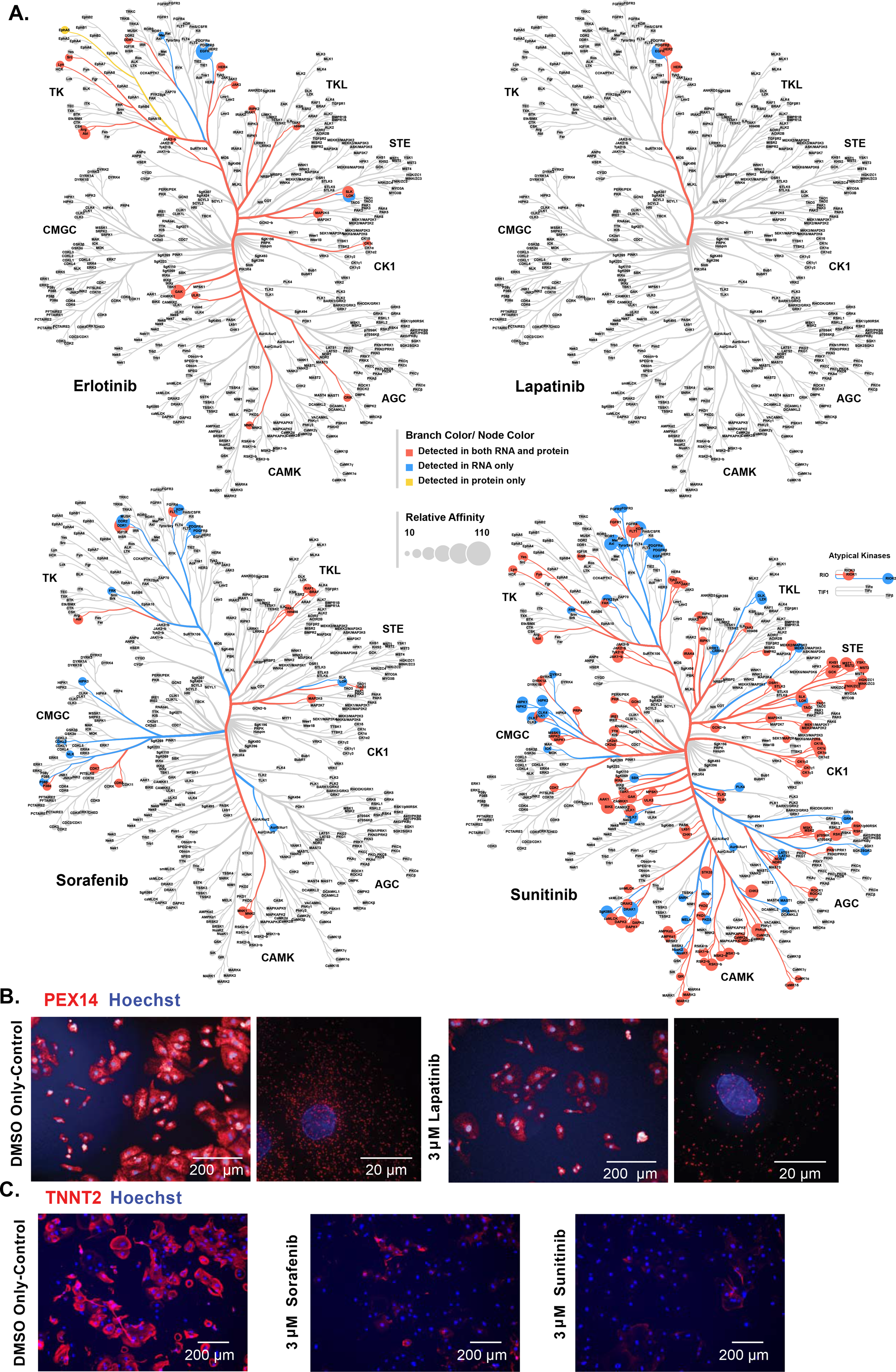
Protein expression changes confirmed by immunofluorescence. (**A**) Kinase targets for each TKI expressed in Cor.4U hiPSC-CMs based on RNA and protein expression data. Target data was obtained from published KINOMEscan profiles and the size of each node represents relative affinity (larger nodes correspond to higher affinity). (**B**) Staining of Peroxisomal Biogenesis Factor 14 (PEX14) in hiPSC-CMs treated with DMSO alone or with 3 μM Lapatinib for 3 days. Low and high magnifications are shown; scale is 200 μM or 20 μM as indicated per field. (**C**) Staining for cardiac isoforms of Troponin T (TNNT2) in hiPSC-CMs treated with a DMSO alone or with 3 μM Sunitinib or Sorafenib at for 4 days. Experiments in Panels A and B were repeated three times and images shown are representative of a single experiment. (See also Figure S5).

Many more gene transcripts were detected by RNA-Seq than proteins by mass spectrometry (across all conditions, ~12,000 coding genes vs. ~7,700 proteins). To compare the two datasets we focused on differentially expressed genes and proteins (DEGs and DEPs) among the 6,151 detected by both assays. PCA of these DEG/DEPs revealed broadly consistent changes (Supplementary Fig. 5A), even though the number of overlapping differentially expressed genes and proteins was quite low. For example, among 753 DEGs and 562 DEPs for cells exposed to 3 μM Sunitinib for 72 hr, we found only 18 overlaps, and among 1412 DEGs and 490 DEPs for cells treated with 3 μM Sorafenib for 72 hr, we found only 98 (Supplementary Fig. 5B). Among the genes and proteins that did overlap, however, the Pearson correlation coefficient for fold-change values was high (r = 0.7 to 1 depending on the drug, Supplementary Fig. 5B). Moreover, GO terms associated with mitochondrial biology, metabolism and sarcomere function were enriched for the overlapping DEGs and DEPs. In several cases this was reflected at the level of individual pathways. For example, DEGs and DEPs mapped to multiple successive steps in cholesterol biosynthesis upregulated by exposure to Lapatinib (Supplementary Fig. 5C) and to multiple components of sarcomeres down-regulated by exposure to Sorafenib (Supplementary Fig. 5D). Coordinated regulation of multiple biosynthetic steps in a pathway or contractile proteins in a common cellular structure are likely to be interesting for follow-up studies aimed at better understanding regulation of the specialized metabolism and contractility of cardiac cells.

### Functional analysis of TKI-induced changes in hiPSC-CM

To confirm the biological significance of TKI-induced changes in gene and protein expression in hiPSC-CMs, we performed functional analyses. First, we looked at proteins involved in peroxisomes and their biogenesis, 18 of which were downregulated 2-fold or more following exposure to 3 μM Lapatinib for 72 hr. This change is detected in proteomic but not RNA-Seq data, implying post-transcriptional regulation of peroxisome protein abundance. When we performed immunofluorescence with antibodies against one of these proteins (PEX14) we found that the number of peroxisomes was reduced ~8-fold by Lapatinib treatment (Fig. 5B). Cardiac cells are particularly dependent on the oxidation of fatty acids for metabolic energy, and much of this oxidation takes place in peroxisomes (van der Vusse et al., 1992); thus, dramatic reductions in peroxisomes in hiPSC-CMs are likely to impose a substantial metabolic burden. Sunitinib and Sorafenib also reduced the expression of Troponin T2, an important component of sarcomeres, ~80-fold in proteomic data as compared to control cells (Fig. 5C) consistent with an observed reduction in spontaneous beating (Supplementary Fig. 1B). Thus, two striking molecular changes associated with TKI exposure appear to have physiological consequences at the levels of organelle structure.

Analysis of TKI-induced metabolic changes focused on Sorafenib, which downregulated genes and proteins involved in mitochondrial electron transport (Fig. 3). Sorafenib has previously been shown to inhibit mitochondrial respiration in tumor cells, immortalized H9c2 rat myocardial cells, and mouse embryonic fibroblasts (Will et al., 2008; Zhang et al., 2017). The effects of Sorafenib on hiPSC-CMs were determined by assaying mitochondrial oxygen consumption rate (OCR on an Agilent Seahorse XF96). Over a 12 min time-course, basal respiratory capacity is measured, followed by treatment with oligomycin A, an ATP synthase inhibitor, to calculate ATP-coupled respiration and then treatment with the mitochondrial uncoupler FCCP to measure maximal respiratory capacity. Finally, Rotenone and Antimycin A (complex I and III inhibitors) were added to measure the rate of non-mitochondrial respiration, which defines the baseline for all calculations. Treatment of hiPSC-CMs with Sorafenib for 48 hr caused a dramatic reduction in mitochondrial function (Fig. 6B, compare red and green lines): basal respiration was reduced 9-fold, ATP-coupled respiration reduced 23-fold, and maximal respiratory capacity was 45-fold lower as compared to control cells (Fig. 6C). Qualitatively similar data were obtained in a second hiPSC-CM line: in this case, Sorafenib reduced basal respiration 3-fold, ATP-coupled respiration 13-fold and maximal respiratory capacity 19-fold (Supplementary Fig. 6).

**Figure 6:**
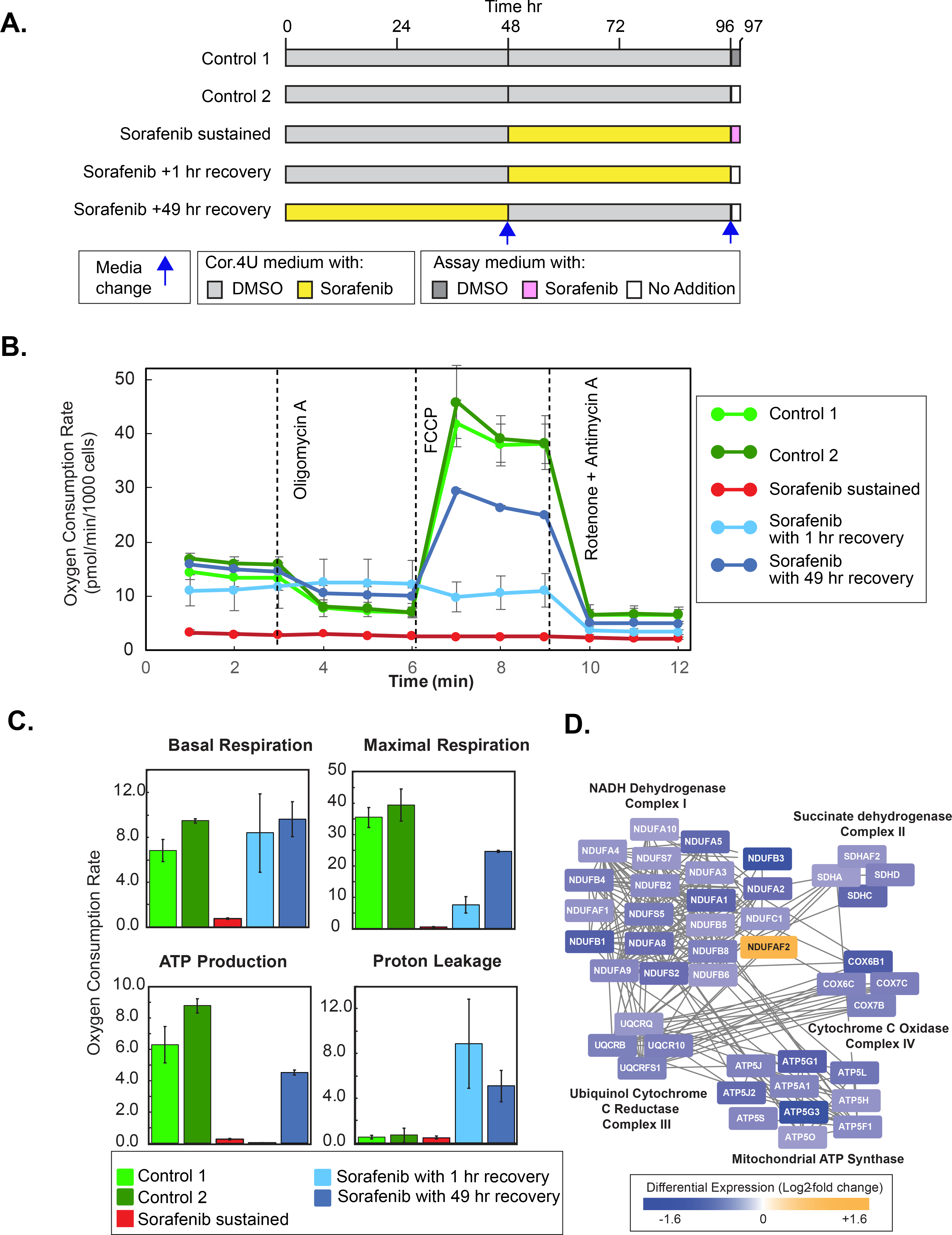
Effect of Sorafenib on mitochondrial respiration in hiPSC-CMs. **(A)** Design of experiments testing the effect of Sorafenib at 6 μM on mitochondrial respiration following different times of recovery. The two controls differ in whether or not DMSO was present in the assay medium. **(B)** Oxygen consumption rate (OCR) in live hiPSC-CMs was measured at baseline and following successive addition of Oligomycin A, FCCP and Rotenone plus Antimycin A (see text for details). Conditions correspond to those described in Panel A. The experiment was repeated twice with 6 replicates per condition per experiment; data were the average from these two experiments. Error bars are standard error with n = 2. **(C)** Metabolic parameters derived from analysis of the OCR data shown in Panel B. Error bars are standard error with n = 2. **(D)** Genes involved in Oxidative Phosphorylation that were down-regulated as measured by RNA-Seq in cells exposed to 10 μM Sorafenib for 24 hr. See Supplementary Figure 6 for confirmation on a second hiPSC cell line.

Changes in mitochondrial function likely reflect reduced expression of multiple subunits of oxidative phosphorylation complexes detected by RNA-Seq (Fig. 6D). To determine whether these changes were reversible, respiration was measured in hiPSC-CMs exposed to 6 μM Sorafenib for 48 hr, followed by a recovery period of 1 hr or 49 hr in which cells were grown in normal media lacking drug (Fig. 6B, blue lines). The study was designed to control for total time in culture, for the duration of Sorafenib exposure, and for the switch (for 1 hr) from Cor.4U growth medium to the Seahorse measurement medium (see Fig. 6A and STAR Methods). We found that 1 hr recovery was sufficient to restore basal respiration but not normal ATP production (as evidenced by the lack of an effect upon oligomycin addition), which was restored only after 49 hr of recovery. However, after 49 hr of recovery, maximal respiratory capacity was still lower than in control cells. Thus, the effects of Sorafenib on mitochondrial function are largely, but not fully, reversible following drug withdrawal. The different time scales for recovery of basal respiration and ATP production suggest that some metabolic processes are directly and reversibly inhibited by Sorafenib (potentially including ATPases other than kinases) and that proteins down-regulated at a transcriptional level take longer to regain function.

### Exposing hiPSC-CMs to Sorafenib results in increased glycolysis as an adaptive response

Glycolysis is an alternative means of producing ATP when mitochondrial function is impaired. Sorafenib treatment of hiPSC-CM was observed to cause an increase in the extracellular acidification rate (ECAR), an indirect measure of glycolysis (Fig. 7A). We detected a decrease in glucose and an increase in lactate in the media of cells treated with Sorafenib as compared to control cells. In addition, hydrophilic interaction liquid chromatography tandem mass spectrometry (HILIC-MS) revealed multiple changes in metabolite levels consistent with a glycolytic shift (see Fig. 7B and Supplementary Table 8) (Spinelli et al., 2017). Thus, hiPSC-CMs compensate for inhibition of mitochondrial respiration by upregulating glycolysis, a shift that is also observed in the adult heart when cells are starved for oxygen.

**Figure 7:**
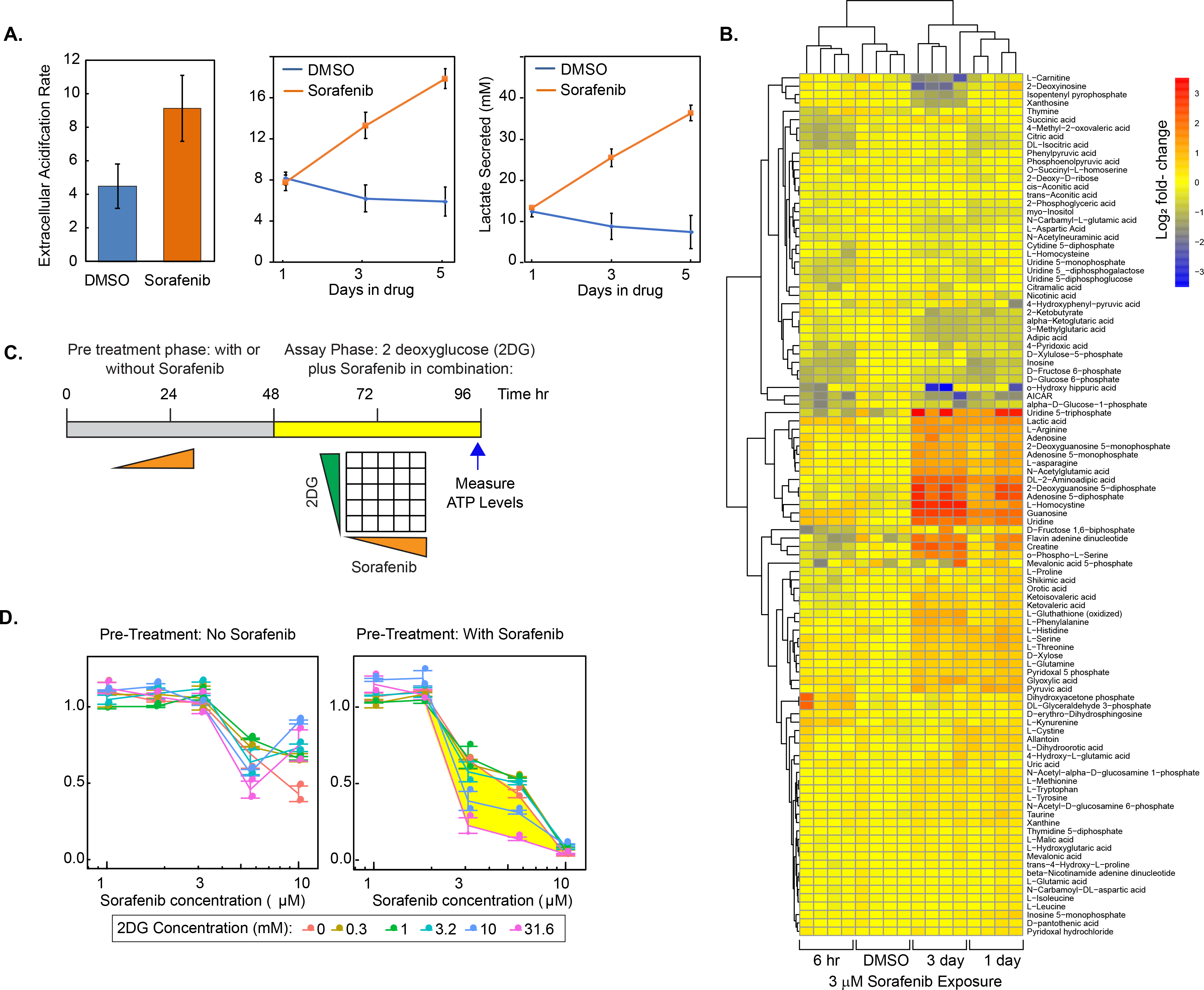
Induction of glycolysis by Sorafenib in hiPSC-CMs. (**A**) Left panel: Extracellular acidification rate (ECAR) measured for hiPSC-CMs cells treated with a DMSO alone or 3 μM Sorafenib for 2 days. Error bars show the standard deviation of one experiment with 6 biological replicates per condition. Data are a representative of study performed three times. Y-axis units are mpH/min/1000 cells. Right panels: Levels of glucose and lactate measured in spent media collected from cells treated with a DMSO alone or 3 μM Sorafenib for 1-5 days. Error bars show the standard error of three independent experiments. (**B**) Metabolic profiling of hiPSC-CMs by hydrophilic interaction liquid chromatography tandem mass spectrometry following exposure of cells to 3μM Sorafenib for 6 hours, 1 day and 3 days; four repeats are shown for each condition. Color scale corresponds to log2-based fold-change in metabolite levels. (**C**) Design and (**D**) data for a co-drugging experiment examining the impact of Sorafenib-induced glycolysis on sensitivity to Sorafenib, as measured using the CellTiter-Glo^®^ luminescence (CTG) assay. In the “pre-treatment phase,” cells were exposed to 0-10 μM Sorafenib for 48 hr and then shifted to an “assay phase” with 0-32 mM 2-deoxy-D-glucose (2DG) and 0-10 μM Sorafenib for 48 hr prior to CTG assay. Data were normalized to luminescence values of cells treated with 2DG alone at the specified doses (see Supplementary Fig. 7A). Each dot represents the average of 3-8 replicates in each of two independent experiments and error bars are the standard error of two independent experiments.

To determine the impact of a glycolytic shift on hiPSC-CM viability, cells were treated with DMSO or Sorafenib at a range of doses (1-10 μM) for 48 hr to induce the glycolytic phenotype, and then with 2-deoxy-D-glucose (2DG; a glucose analog that competitively inhibits hexokinase) in combination with Sorafenib for another 48 hr (Fig. 7C). We then measured ATP levels using the CellTiter-Glo assay (Fig. 7D). As controls, we measured the effect of 2DG treatment alone and of 2DG plus Sorafenib in the absence of Sorafenib pre-treatment. hiPSC-CMs culture media contains ~25mM glucose and we therefore tested 2DG over a 0-32 mM range. As expected, 2DG alone inhibited metabolism in a dose-dependent manner, reducing ATP to ~60% of control levels after 48 hr in the presence of 32 mM 2DG (Supplementary Fig. 7); co-treatment of cells with 2DG unexpectedly reduced the impact of 10 μM Sorafenib on ATP levels for reasons that are unknown, but hiPSC-CMs rapidly lose viability in 10 μM Sorafenib (Fig. 7D). To determine whether the shift to glycolysis at lower Sorafenib doses was physiologically important, we divided the effect on ATP levels of Sorafenib plus 2DG by the effect of 2DG alone to derive a dose response of ATP levels to Sorafenib. When 2DG and Sorafenib were added together across a range of doses in the absence of pretreatment, inhibition of aerobic glycolysis by 2DG minimally affected response to Sorafenib (Fig. 7D). However, when cells were pre-treated with 3 or 6 μM Sorafenib for 48 hr, 2DG (particularly 10 and 31.6 mM) made hiPSC-CM cells substantially more sensitive to Sorafenib treatment (in yellow, Fig. 7D). Thus, the glycolytic shift observed in Sorafenib-exposed hiPSC-CMs is an adaptive response to drug necessary for cardiomyocytes to preserve their metabolic activity.

## DISCUSSION

In this paper we profile changes in gene and protein expression and the physiology of hiPSC-CMs exposed to four TKIs used to treat solid tumors. The long-term goal of such an analysis is understanding and eventually mitigating drug-induced cardiotoxicity. hiPSC-CMs are a renewable resource for studying cardiac molecular biology, well suited to counter-screening in drug development and much more experimentally tractable than whole animals. However, hiPSC-CMs have some limitations, including functional but not fully aligned sarcomeres, molecular properties midway between fetal and adult cells and, in this study, growth as monolayers on plastic – a biologically unrealistic condition used because it facilitates molecular profiling.

We find that many of the kinases targeted by TKIs are expressed in cardiac cells. These kinases, and the signaling pathways in which they function, are known from other studies to be important for cardiomyocyte physiology (Chen et al., 2008). Inhibition of signaling kinases results in cell cycle arrest and death of cancer cells, and similar activities are postulated to cause TKI-induced cardiotoxicity (Cheng and Force, 2010; Force et al., 2007). However, we find that TKIs induce wide-spread reprogramming of gene expression in hiPSC-CMs at lower drug doses, conditions in which little or no cell death is observed. Systematic profiling is consistent with previous studies in demonstrating Sorafenib-mediated up-regulation of ER stress genes (e.g. DDIT3, ATF3 and CHAC1; (Dixon et al., 2014) and changes in the expression of VEGF proteins (e.g. KDR and VEGFC; (Sharma et al., 2017)). Lapatinib is also known to cause widespread dysregulation of cholesterol biosynthetic enzymes (Necela et al., 2017), as observed here for hiPSC-CMs. We also find that many cell cycle genes are down-regulated when hiPSC-CMs are exposed to TKIs. This is surprising since adult cardiomyocytes are post-mitotic, but we find that a substantial fraction of Cor.4U hiPSC-CMs are positive for the S-phase marker phosphorylated Rb (p-pRb), even those cells positive for the differentiation markers Mlc-2v and Mlc-2a. The fraction of p-pRb positive cells was reduced by TKI exposure, consistent with the known anti-mitogenic effects of these drugs (Supplementary Figure 2C), and with the results from RNASeq. We speculate that TKI-induced changes in levels of cell cycle genes are an artefact of the relative immaturity of hiPSC-CMs and not relevant to adult tissues. However, pRb expression has been observed in adult cardiomyocytes during cardiac hypertrophy (Hinrichsen et al., 2008; Wohlschlaeger et al., 2010; Yuan et al., 2016) and following hypoxia (Sogawa et al., 1998). It is therefore possible that changes in the expression of cell cycle genes are part of a cardiomyocyte stress response. Distinguishing between these possibilities will require comparison of hiPSC-CMs and primary adult cells.

We find that GO processes corresponding to metabolism, biosynthesis, and remodeling of the extracellular environment are also dysregulated by multiple TKIs. For example, Lapatinib exposure downregulates multiple peroxisomal proteins and dramatically reduces peroxisome number (Figure 5); Lapatinib also induces expression of cholesterol biosynthetic enzymes. Since peroxisomes play an important role in cholesterol biosynthesis and homeostasis (Faust and Kovacs, 2014), the two biological pathways affected by Lapatinib may be connected functionally: upregulation of cholesterol pathway enzymes may be a response to a loss of peroxisomal biosynthetic capacity.

Drug dose and time of exposure are key pharmacological variables often ignored in genome-wide profiling studies because each condition is expensive to analyze. We used simple functional assays to explore drug dose and exposure times and identify conditions that elicit detectable cellular phenotypes but not cell killing. For Lapatinib, Erlotinib, and Sorafenib optimal *in vitro* concentrations were similar to or below C_max_ values in humans, but for Sunitinib they were 10-fold higher. Why this is true is not clear, but drug doses needed to elicit therapeutic proxy effects in cultured cells (e.g. cytostasis or cytotoxicity) can also differ from those *in vivo*. Based these exploratory studies, molecular profiling was performed using a relatively compact set of six conditions spanning dose and time. For some drugs and molecular processes (as defined by enriched GO terms), total exposure was the key variable: enriched genes under high-dose and short time conditions were highly correlated with those at lower dose and longer time including the metabolism-linked changes described above. In these cases it seems likely that *in vitro* studies spanning a period of days will be relevant to understanding human exposure spanning a period of weeks to months.

The data in this paper represent a resource for generating network-level hypotheses about drug-induced cardiotoxicity. We have mined only a small subset of the gene enrichment and GO data, and validated only peroxisomal, sarcomere and mitochondrial phenotypes. Deeper analysis of these phenomena is clearly warranted. To this end, all data have been annotated to NIH LINCS standards and deposited in public repositories (Synapse syn13856689, PRIDE PXD012043 and GEO GSE114686). Molecular profiles were derived from multiple independently differentiated lots of Cor.4U hiPSC-CMs over a period of ~18 months, potentially introducing batch effects. However, GSEA results were well-correlated across lots, suggesting that the dataset does not have serious confounders. Because Cor.4U cells are commercially available, it should be straightforward for others to add to the dataset and test specific hypotheses experimentally.

The metabolic defect we have studied most carefully is Sorafenib-dependent inhibition of mitochondrial ATP production and a concomitant increase in glycolysis. The adult myocardium is highly metabolically active and 95% of its energy derives from oxidative phosphorylation in mitochondria fueled by fatty acyl-coenzyme A. The effect of Sorafenib on mitochondria is reversible upon drug withdrawal, as is Sorafenib cardiotoxicity in human patients (Uraizee et al., 2011), and probably reflects the normal ability of cardiomyocytes to switch energy source as a function of developmental stage and oxygen levels (Breckenridge et al., 2013). Induction of glycolysis reduces the toxicity of Sorafenib, presumably by providing an alternative source of ATP. However, increased reliance on glycolysis is a characteristic of cardiac hypertrophy, myocardial ischemia and cardiac failure (Doenst et al., 2013; Rosano et al., 2008; Ventura-Clapier et al., 2004). Sorafenib and other TKIs have also been observed to reduce blood glucose levels (Agostino et al., 2011), potentially enhancing the adverse consequences of reliance on glycolysis for energy. Thus, in the short-term, a switch in Sorafenib-treated hiPSC-CMs from oxidative phosphorylation to glycolysis is a form of adaptive drug resistance, but in the longer term (i.e. in patients) such a switch may be toxic. Thus, we propose that Sorafenib cardiotoxicity might be a consequence of metabolic remodeling rather than by inhibition of signal transduction, as previously suggested (Force et al., 2007).

Adaptive drug resistance has most extensively been studied in tumor cells and combination therapies are being developed to block adaptation and increase tumor cell killing. However, in the case of cardiomyocytes, blocking adaptation or decreasing blood glucose levels by indirect means is expected to increase the adverse effects of Sorafenib. Thus, our data suggest that managing drug-induced cardiotoxicity should focus not just on acute on-target effects of drug exposure, but also adaptive and maladaptive physiological responses.

## Supporting information

Key Resources Table

## ACKNOWLEDGMENTS

We thank J. Lin, R. Jiang, A. Jenney, S. Gygi, and scientists from Qiagen Lifesciences and Axiogenesis. The research was supported by FDA contract HHSF223201400052C and NIH grants U54-CA225088 and U54-HL127365 to PKS, and AHA fellowship 15POST25230014 and NIH F32 HL14223 to HW. The manuscript is dedicated to Darrell Abernethy MD PhD, an FDA Associate Director who inspired our research until his untimely death.

## Author Contributions

HW, LM and PKS conceived the study. HW performed experiments except that: SAB performed RNA-Seq, RAE and ARE mass spectrometry, NRH and AER metabolite profiling (supervised by MCH), CAJ immunofluorescence, RPS and ACP data analysis. KMH assisted with edgeR and goseq. HW, RPS and PKS wrote the paper.

## STAR METHODS

### Contact for Reagent and Resource Sharing

Further information and requests for resources and reagents should be directed to and will be fulfilled by the Lead Contact, Peter Sorger, at.

## EXPERIMENTAL MODEL AND SUBJECT DETAILS

### Human induced pluripotent stem cell-derived cardiomyocytes (hiPSC-CMs)

Cor.4U hiPSC-CMs were purchased as differentiated cardiac myocytes derived from a female or a male hiPSC line by the manufacturer (Ncardia; previously Axiogenesis; https://www.ncardia.com). The female hiPSC-CMs has been commercialized, but the male hiPSC-CMs was an internal line shared with us by the manufacturer. hiPSC-CMs were maintained in the manufacturer’s media: *growth media* (Cor.4U Complete Culture Medium which contains 10% FBS and other essential nutrients, Catalog number Ax-M-HC250 from Ncardia) or *minimal media* (BMCC Serum-Free Culture Medium, Catalog Number Ax-M-BMCC250 from Ncardia, supplemented with 1% fetal bovine serum from Thermo Fisher Scientific) under 5% CO2 at 37 °C in a Biosafety Level 2 facility.

## METHOD DETAILS

### Culture of hiPSC-CMs

Cryo-preserved hiPSC-CMs were thawed into 75 cm^2^ flasks following the manufacture’s protocol, cultured for 3 days, then reseeded into multi-well plates to remove dead cells caused by thawing. Cells were initially cultured in growth media and then switched to minimal media 1.5 days prior to drug treatment. hiPSC-CMs were maintained in minimal media during drug treatment to minimize interference from serum with the drug-induced effects. During culture, media were exchanged every other day.

### Drug treatment

Further information on the Tyrosine Kinase Inhibitors (TKIs) used in this study, Erlotinib, Lapatinib, Sorafenib and Sunitinib, is provided in the “Key Resources Table.” All drugs were assayed for purity and concentration by NMR in-house after which they were dissolved in DMSO, aliquoted and stored at −20°C. 2-deoxy-D-glucose (2DG) was dissolved in water. Drug treatment of cells in 384-well plates was performed as shown in Figure 1A or randomized across the plate using an HP D300 digital dispenser (Hewlett-Packard). Master drug stocks were stored at 10 mM in DMSO for TKIs and 2 M in water for 2DG with 0.03% Tween 20. Tween was added to facilitate dispensing of drugs in aqueous solvent by the HP D300 dispenser. Drug treatment of cells cultured in 6-well or 60 mm plates (for omics experiments) was performed by manual pipetting. The overall experimental design for phenotypic, transcriptomic and proteomic experiments was shown in Supplementary Fig. 1C.

### Live cell high-content imaging of mitochondrial membrane potential

Following pre-culture, hiPSC-CMs cells were re-seeded at ~4000 cells/well in optically clear 384-well plates (Corning) in minimal media and treated with drugs at a range of doses (0.1 to 10 μM) and times (6 hours and 1, 3, 5 and 7 days; see Supplementary Fig. 1C) in 2-3 biological replicates. At each time point, cells were stained for 30 min in a cell culture incubator (5% CO_2_ and 37°C) with a mixture of Tetramethylrhodamine ethyl ester perchlorate (TMRE, Sigma) at 1 nM, Calcein (Life Technologies) at 0.1 μM, MitoTracker Deep Red (Life Technologies) at 50 nM, and Hoechst 33342 (Life Technologies) at 162 nM. Cells were imaged using an Operetta High-Content Imaging System equipped with a live cell chamber (Perkin Elmer).

### CellTiter-Glo Assays

ATP content (data shown in Fig. 1B and 7C in the main text) was measured using the CellTiter-Glo Luminescent Cell Viability Assay (Promega) following the manufacturer’s protocol; luminescence was quantified using a Synergy H1 plate reader (BioTek Instruments). Assay conditions were optimized using ATP standards (0.01-5 μM) and all luminescence readings were found to fall in a linear range for the CellTiter-Glo assay.

### Calcium flux assay

Following pre-culture, hiPSC-CMs cells were seeded in black 384-well plates with a transparent bottom (Corning) in minimal media and treated with drugs at a range of doses (0.1 to 10 μM) and times (6 hours and 1, 2, 3 and 6 days) in 3 biological replicates. At each time point, cells were stained for 30 min with an EarlyTox Cardiotoxicity Kit (Molecular Devices), which contains a less toxic modified calcium dye than traditional dyes, in a cell culture incubator (5% CO_2_ and 37°C). Fluorescence intensity was recorded using a FDSS 7000 System (Hamamatsu) at 37°C. Beats per minute were calculated using the waveform add-on software for the FDSS 7000 device (Hamamatsu).

### Total RNA sequencing

RNA-Seq data are available from GEO with the accession number GSE114686. hiPSC-CMs were pre-cultured as in Fig. 1A. Cells were then treated with one of four TKIs at three doses (1, 3 and 10 μM) for 24 hours or at a single dose (3 μM) for four different times (6 hours, 1, 3 and 7 days), or with a DMSO as a control (DMSO-only data were collected at 6 hours, 1, 3 and 7 days to monitor changes in Cor.4U cells in the absence of drugs). A different lot of Cor.4U cells was used for each of three independent RNA-Seq experiments. Drug-containing media was refreshed every other day. Cell lysates from the 6-hour treatment were collected 21 hours after the last media change; cell lysates of 1, 3 and 7 day treatments were collected 39 hours after the last media change (see Supplementary Figure 2F).

Cell lysate for RNA isolation was prepared as follows. Cells cultured and drug-treated in 6-well plates were washed with PBS, harvested in RLT buffer from the RNeasy Mini Kit (Qiagen), and frozen at −80°C until further processing. At the time of RNA purification, samples were thawed at room temperature and processed on QIAshredder (Qiagen) columns to ensure complete lysis. Total RNA purification was performed following the manufacturer’s protocol for the RNeasy Mini Kit (Qiagen), including the manufacturer recommended 30 minute on-column DNase digestion step. Total RNA concentrations were measured by Nanodrop assay and quality was assessed on a subset of samples using a Bioanalyzer (Agilent); all samples had RIN scores of > 9.0.

RNA sequencing library preparation was performed using the High Throughput TruSeq Stranded Total RNA Library Prep Kit with Ribo-Zero gold (Illumina) following the manufacturer’s protocol. 500 ng of RNA was input for each sample and supplemented with 1 μl of 1:100 diluted ERCC spike in mix A (Ambion). Libraries were amplified for 15 cycles at the final step of library preparation. Amplified libraries were quantified using Qubit dsDNA HS assay (Thermo Fisher Scientific) and library size and quality was checked on a subset of samples by Bioanalyzer (Agilent). The resulting libraries were composed of cDNA fragments of 330 base pairs on average. Samples were pooled at equimolar concentrations and sequenced using HiSeq 2500 V4 (Illumina) at the Bauer Core Facility (Harvard University).

### Tandem Mass Tag Mass Spectrometry

Proteomics raw data are available from PRIDE under project accession number PXD012043. Two independent drug treatment experiments were performed. In the first experiment, hiPSC-CMs were cultured in 60 mm plates, then treated for 3 days with 3 μM of each of the four TKIs (Erlotinib, Lapatinib, Sorafenib and Sunitinib) or with as a DMSO vehicle-only control, in duplicate (10 samples in total). In the second experiment, a different lot of hiPSC-CMs were cultured and treated with each of the four TKIs at 3 μM or DMSO, and samples were collected at both day 1 and day 3. This resulted in 30 samples that were processed in three 10-plex TMT-labelling experiments. 10 samples from each time point comprised one 10-plex TMT run (see Supplementary Fig. 4A).

Each 10-plex TMT sample as prepared as follows. Cells cultured in 60 mm plates were washed 3 times with PBS, lysed and scraped in 0.5 mL of lysis buffer (2% SDS, 150 mM NaCl, 50 mM Tris, pH 8.5) containing protease and phosphatase inhibitors (Roche). The cell lysate was collected and vortexed, then passed through a QIAshredder (Qiagen) column to ensure complete lysis. The protein concentration of each sample was determined using 30 μL of cell lysate measured by BCA (Thermo Fisher Scientific). The remaining cell lysate was reduced with 5 mM DTT and alkylated with 15 mM iodoacetamide (Sigma-Aldrich). For each sample, 0.15 mg of protein was aliquoted into separate tubes and precipitated by adding 3x volumes of chloroform and 2x volumes of methanol. The pellet was recovered by centrifugation and washed with cold methanol 2 times and re-solubilized in 8M urea with 20 mM EPPS (Sigma-Aldrich), pH 8.5. The sample was then diluted into 4M urea with 20 mM EPPS and digested using Lys-C (Wako Chemicals) overnight at room temperature. The sample was further diluted to a final concentration of 1.5 M urea with 20 mM EPPS and digested with Trypsin (Promega) for 6 hours at 37 °C. 60 μg of each sample was then dissolved into 10% (v/v) acetonitrile (ACN) and labeled 2:1 (TMT:Peptide ratio) with TMT-10 reagent (Thermo Fisher Scientific). The reaction was quenched with hydroxylamine (0.5% final volume, Sigma-Aldrich). The ten samples per experiment were then combined, acidified by adding formic acid (FA, Thermo Fisher Scientific) to 2% final volume, and desalted using a C18 Sep-Pak (Waters). The combined sample was fractionated using basic pH reversed phase chromatography on an Agilent 1200 HPLC equipped with a UV-DAD detector and fraction collection system. A total of 12 fractions were desalted using the C18 StageTip procedure. Each fraction was loaded onto a column (35 cm long x 100 μm inner diameter) packed with 1.8 μm beads (Sepax) and separated using a 3-hour gradient from 8% Buffer B and 92% Buffer A to 27% Buffer B and 73% Buffer A (Buffer B: 99% ACN and 1% FA. Buffer A: 96% water, 3% ACN and 1% FA) using an Easy 1000 nano-LC (Thermo Fisher Scientific). All MS analyses were performed on an Orbitrap Fusion Lumos mass spectrometer (Thermo Fisher Scientific) using a multi-notch MS3 method.

### Assaying metabolites in conditioned (spent) media

Spent media was recovered from the three independent total RNA-Seq experiments at day 1, 3 and 5. Spent media samples were always collected at 39 hours after regular media changes and were store at −80°C until measurement. Glucose and Lactate concentrations (in mM) were measured in undiluted media using the BioProfile Flex Analyzer (Nova Biomedical). Culture media was incubated under the same conditions and for the same amount of time as Cor.4U cell cultures to serve as a control for measurement of metabolite concentrations in the absence of cells.

### Metabolomics of cell extracts

Cor.4U cells were treated with either vehicle-only control (0.1% DMSO) for 3 days or Sorafenib at 3 μM for 6 hours, 1 day and 3 days. Reverse scheduling of treatment was used so that all samples were collected at the same time and cells were in culture for the same total amount of time. During metabolite collection, cells were washed with 1 mL ice-cold PBS, aspirated, incubated in 1 mL of 80% HPLC-grade Methanol (for a 10 cm dish) for 10 minutes at −80 °C and scraped on dry ice. Lysates were centrifuged at 10,000 x g for 10 minutes and supernatants were used for the metabolome measurements.

Metabolites from biological samples were resolved using reverse phase ion-paired chromatography on an Agilent 1290 Infinity II Series LC and detected by MRM in negative ion mode on an Agilent 6470 series triple quadrupole mass spectrometer. The LC-MS/MS targeted MRM method used for metabolomics analysis was developed and implemented by Agilent Technologies (https://www.agilent.com/en/products/mass-spectrometry/analyzers-databases-libraries/life-sciences/metabolomics-dmrm-database-method). In brief, samples were dried down in a speed-vac at room temperature, resuspended in Buffer A (97% H2O, 3% Methanol, 10 mM Tributylamine, 15 mM Glacial Acetic Acid, pH 5.5), and then 10 uL was injected over a ZORBAX Extend-C18, 2.1 × 150 mm, 1.8 μm column (Agilent) equilibrated in Buffer A at a flow rate of 0.25 mL/minute. The column was developed with the following method: 2.5 minutes with 0% Buffer B (10 mM Tributylamine, 15 mM Glacial Acetic Acid in 100% Methanol), linear gradient of 0-20% Buffer B over 5 minutes, linear gradient of 20-45% Buffer B over 5.5 minutes, linear gradient of 45-99% Buffer B over 7 minutes, and finally 4 minutes with 99% Buffer B. Between runs, the column was cleaned with an acetonitrile backwash and then the column was re-equilibrated for 8 minutes in 100% Buffer A. Samples were ionized in negation ion mode using an Agilent Jet Stream Source with the following MS source parameters: nebulizer = 45 psi, capillary voltage = 2000V, nozzle voltage = 500 V, sheath gas temperature = 325°C, sheath gas flow = 12 L/minute, gas flow = 13 L/minute, and gas temperature = 150°C.

### Mitochondrial respiration assay

hiPSC-CMs were seeded into Seahorse XF96 cell culture plates (Agilent) following pre-culture and treatment with drugs as described in the main text. About 2.5 hours prior to the assay, plates were switched to a 37°C non-CO_2_ incubator for 1.5 hours. One hour prior to the assay, cells were changed to the Seahorse measurement medium, which is made of the Seahorse Basal DMEM medium with 5 mM glucose, 2 mM L-glutamine, and 1 mM sodium pyruvate. The Seahorse Basal DMEM medium differs from the regular DMEM in that it is free of bicarbonate, low in buffer capacity, and contains low phenol red (according to the manufacturer). Oxygen consumption rates (OCR) and extracellular acidification rates (ECAR) were measured in live hiPSC-CMs cells under a sequential drug treatment involving Oligomycin A (1 μM, an inhibitor of ATPase), Carbonyl cyanide 4-(trifluoromethoxy) phenylhydrazone (FCCP, 1 μM, a mitochondrial uncoupler), and a combination of Rotenone (0.5 μM, an inhibitor of mitochondrial complex I) and Antimycin A (0.5 μM, an inhibitor of mitochondrial complex III), using an XF-96 Extracellular Flux Analyzer (Agilent). For the data shown in the Main Text Figure 6 and Supplementary Figure 6, hiPSC-CMs were treated with DMSO at 0.1% (vehicle-only control) or Sorafenib at 3 μM for 2 days in minimal media. When the media were replaced with Seahorse measurement media one hour prior to the assay, drug was removed. For the metabolism data shown in Main Text Figure 7, hiPSC-CMs were maintained in culture for a total of 4 days and exposed to Sorafenib at 6 μM for the first or the last 2 days, which corresponds to recovery from Sorafenib treatment for either 49 hours or 1 hour prior to the assay (see Results for details). During the culture time that Sorafenib was not present, DMSO at 0.1% was added. The vehicle-only control was treatment with DMSO (0.1%) for 4 days. When media were replaced to the Seahorse measurement media one hour prior to the assay, drugs were either refreshed or removed depending on the condition. These experiments were done with 6 replicates per condition at two independent times.

### Immunofluorescence

Cultured, treated cells were fixed for 20 min using 4% Paraformaldehyde (Electron Microscopy Sciences) at room temperature and washed with PBS at least three times before staining. Cells were permeabilized with 0.1% TritonX100 in PBS for 15min, and blocked with Odyssey blocking buffer (Li-Cor) for 1 hour. Samples were then incubated overnight at 4°C with the primary antibodies at the optimal dilution, washed with PBS three times, and incubated with Alexa Fluor-based secondary antibodies (Thermo Fisher Scientific) and DAPI (Sigma-Aldrich) for 1 hour at room temperature (see Key Resources Tables for the identities of all antibodies used in this study). Cells on slides were mounted with Prolong Gold mounting media (no DAPI, Thermo Fisher Scientific). Samples were imaged using the Cytell Imaging system (GE Healthcare Life Sciences), the DeltaVision Elite Imaging System 7 (GE Healthcare Life Sciences), or the Operetta High-Content Imaging System (Perkin Elmer). The TNNT2 staining in Figure 5 was based on the mouse TNNT2 antibody and that of Figure 1B and Supplementary Figure 1 was based on the Rabbit TNNT2 antibody. For co-staining of pRb and Mlc2v and Mlc2a in Supplementary Figure 2C, we used the CyCIF method (Cyclic Immunofluorescence) developed in the Sorger lab. Cells were first stained with the anti-Mlc2v antibody and a secondary antibody labeled with Alexa Fluor 488, following the standard immunofluorescence staining procedures described above. Then, samples were treated with a mixture of 3% H2O2 and 20 mM NaOH in PBS (137 mM NaCl, 2.7 mM KCl, 10 mM Na2HPO4 and 1.8 mM KH2PO4; pH≈9.5) for 1 hr at room temperature to bleach the fluorophore and washed with PBS for three times. A second round of staining was performed using the anti-Mlc2a antibody conjugated with ATTO 647N. Samples were bleached again, washed and stained with the anti-pRb antibody conjugated with Alexa Fluor 555. Samples were imaged at InCell 6000 (GE Healthcare). Primary antibodies used are listed in the “Key Resources Table.”

### Quantitative Real Time-Polymerase Chain Reaction (qRT-PCR)

qRT-PCR was performed in three replicate wells per condition. Cor.4U cells were pre-cultured, plated and treated with each of the four TKIs at 3 μM for 24 and 72 hours, as well as at 10 μM for 24 hours. As a control, cells were also treated with a DMSO vehicle-only control for 24 and 72 hours. A total of 42 RNA samples were isolated using RNeasy Mini Kit (Qiagen) and each sample was found to have a 260/280 ratio >=2.0. RNA samples were shipped to Qiagen Lifesciences Service Core (Frederick, MD) for further processing and performance of qRT-PCR. At Qiagen the samples were reverse transcribed to cDNA using an RT^2^ First Strand Kit (Qiagen). For RT-PCR 90 genes (Supplementary Table 3) were selected based on RNA-Seq data as were four control house-keeping genes (ACTB, GUSB, NONO, RPLP0) using custom RT^2^ profiler arrays and RT² SYBR Green ROX qPCR Mastermix (Qiagen). The QuantStudio 7 Flex Real-Time PCR System (Applied Biosystems) was used to measure Ct values from PCR amplification data (Thermo Fisher Scientific).

## QUANTIFICATION AND STATISTICAL ANALYSIS

### High content imaging analysis

Images acquired as described above were exported to and analyzed in the Columbus Image Data Storage and Analysis System (Perkin Elmer). Calcein staining was used to create single cell masks. Cell masks were filtered based on a cell area > 500 μm^2^, and a nuclear area between 60 and 500 μm^2^ and border cells were removed. All fluorescence intensity was calculated by subtracting the mean fluorescence intensity in cells to that of cell-free regions in the same plate or well. Normalized mitochondrial membrane potential was calculated by (TMRE intensity/MitoTracker Deep Red intensity) normalized to the DMSO control at the same time point.

### RNA-Seq analysis

#### Differential gene expression analysis

Raw RNA-Seq reads were mapped onto human genome hg19 using the STAR aligner packaged in the bcbio-nextgen suite. One sample failed library preparation yielding results for 83 samples. Three of the 83 RNA samples were deemed to be contaminated by ribosomal RNA (rRNA) since their percentage of rRNA reads exceeded 10%. These were removed from the subsequent differential expression analysis.

Based on the count table generated from STAR alignment, we performed differential expression analysis using edgeR. A filtering criterion of mean Count Per Million (CPM) > 1 of each time point in the vehicle-only controls resulted in 13,227 transcripts. Replicate 3 was found to have a consistent batch effect distinct from the other 2 replicates; thus, we normalized for the batch effect by adding a factor for “replicate number” into the experimental design matrix in edgeR. Statistical analysis of differential expression was performed by fitting a negative binomial generalized log-linear model to the read counts and conducting gene-wise statistical tests for a given contrast. Likelihood ratio test (glmLRT) with Benjamini and Hochberg multiple-test correction was used to compare drug-treated samples to the vehicle control at the same time point. Three independently sampled biological replicates, 8567 transcripts were significantly regulated by any one treatment condition with adjusted p≤0.05 and fold-change ≥1.5.

### Principal component analysis (PCA)

Log2-based fold-changes induced in the 13,227 measured transcripts for each drug were calculated using edgeR. PCA was performed based on a data matrix where the columns were the 6 treatment conditions (as shown in Figure 2A) and the rows were composed of log2-fold-changes of four drugs concatenated. After the initial PCA, we rotated the first two principle components (PC1 and PC2) by *θ* =45° using a rotation matrix, 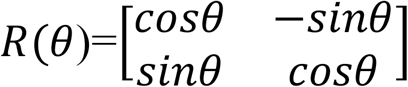, and this resulted in two new PCs: one primarily explained by dose-induced gene expression changes (PC1’) and the other by time-induced changes (PC2’).

### g-means clustering coupled with goseq

Log2-based fold changes of 8567 significantly regulated transcripts were clustered using a g-means algorithm, an abstraction of k-means clustering which optimizes the number of clusters by continuously splitting each cluster until it conforms to a Gaussian distribution. The algorithm was run using a significance level of alpha = 0.001, splitting clusters deterministically based on the largest eigenvector in the old cluster. 16 clusters were required to best fit the data (see Supplementary Table 4). Genes from each cluster were analyzed using goseq and genes in the top 2-3 enriched GO terms were visualized in heat maps.

### Analysis of Doxorubicin data from GEO

Six datasets were collected form the NCBI Gene Expression Omnibus (GEO) database, with accession numbers GSE12260, GSE42177, GSE81448, GSE64476, GSE97642, and GSE40289. Each dataset resulted from measurement of transcriptomic changes in the heart or in isolated cardiomyocytes following treatment with Doxorubicin. Mean differential gene expression across all replicates was calculated using the GEO2R web-based tool, and genes with a p-value <= 0.05 were used for GO term enrichment analysis using the Enrichr web-based tool (http://amp.pharm.mssm.edu/Enrichr/). Additionally, differential expression values for all genes found in each of the six datasets, as well our TKI-treated RNA-seq dataset, were used in a PCA analysis. Each dataset was normalized to ensure a mean of 0 and standard deviation of 1, and the MATLAB built in PCA method was used, using the eigenvalue decomposition method.

### Analysis of Proteomics data

#### Differential expression analysis

Raw data were converted to mzXML and mapped via Sequest version 28 against a concatenated Uniprot database downloaded on 02/04/2014. Variable modifications of oxidized Methionine and over-labeling of TMT on Serine, Threonine and Tyrosine were considered. Linear discriminate analysis was used to distinguish forward and reverse hits, and reverse hits were filtered to an FDR of 1% at the protein level. Shared peptides were collapsed into the minimally sufficient number of proteins using rules of parsimony. Quantitation filters of > 200 sum reporter ion S:N and > 0.7 isolation specificity were incorporated. After converting raw data into values of signal intensity based on these pre-processing steps, data were further normalized so that protein expression values for the 10 samples of each protein sum up to 100 in each 10-plex TMT experiment. The two 10-plex TMT experiments of 72-hour treatment, were combined by calculating the mean of one replicate from each of the two experiments. To confirm the validity of combining the results from the two independent experiments, we performed a principal component analysis. As expected, the four replicates for each condition, spanning both experiments, consistently cluster together (see Supplementary Figure 4A).

One-way ANOVA with Bonferroni correction was conducted for each protein after data combination, and significance was determined based on adjusted p ≤ 0.05 and fold-change ≥1.2. PCA was performed for log2-based fold-changes in protein expression induced by different drug treatment.

### Putative target mapping

Kinase trees shown in Figure 5A were generated using the online tool Coral: http://phanstiel-lab.med.unc.edu/CORAL. For each TKI, the size of each nodes in the free was scaled as log10(1/Kd x 10000) x 20 so that targets associated with a lower Kd value are bigger. Note that two of five putative targets of Lapatinib (https://www.discoverx.com/tools-resources/reference-compound-data) are absent from the corresponding tree because the they are not found in the source data for Coral (Supplementary Table 6).

### k-means clustering coupled with goseq

Similar to the RNA-Seq data analysis, log2-fold-changes of differentially expressed proteins were clustered using k-means clustering with k set to 11. For each cluster, we used the Bioconductor package goseq to perform GO term enrichment and KEGG pathway enrichment analysis.

### qRT-PCR data analysis

Ct values were exported to a text file and analyzed using R. Ct values were normalized to the average Ct values of three of the housekeeping genes (GUSB, NONO, RPLP0) whose expression was determined to be unchanged among the experimental conditions. One-way ANOVA with Bonferroni correction was used for differential expression analysis. Log2-fold changes measured in qRT-PCR were compared with those measured in RNA-Seq.

### Statistical analysis

In Figure 7B, media were collected from 1-3 replicate wells per experiment and from 3 independent experiments. Student’s t test was used to compare drug treatment and control at the same times points (n=3). Significance was determined based on p value ≤ 0.05. In Figure 7D, drug responses were measured with 3-8 replicates per treatment in each of the two independent experiments, and percent cell survival was calculated by normalizing to the treatment of 2DG alone at the specified doses.

## DATA AND SOFTWARE AVAILABILITY

The following are available for download; imaging data are available on request.

- The RNA-seq data have been deposited in GEO (https://www.ncbi.nlm.nih.gov/gds) under ID GSE114686
- The proteomics data have been deposited in the PRIDE Archive (https://www.ebi.ac.uk/pride/archive/simpleSearch) under ID PXD012043
- Synapse (syn13856689): a Wiki page about this project and all associated omics datasets (https://www.synapse.org/#!Synapse:syn13856689)
- Github: Data analysis software (https://github.com/sorgerlab/CardiotoxDataAnalysis)
- Harvard Medical School Library of Integrated Network-based Cellular Signatures (LINCS) center datasets: qRT-PCR data: http://lincs.hms.harvard.edu/db/datasets/20325/; RNA-Seq data: http://lincs.hms.harvard.edu/db/datasets/20324/
- Imaging data are available upon request

## LITERATURE CITED

Agostino, N.M., Chinchilli, V.M., Lynch, C.J., Koszyk-Szewczyk, A., Gingrich, R., Sivik, J., and Drabick, J.J. (2011). Effect of the tyrosine kinase inhibitors (sunitinib, sorafenib, dasatinib, and imatinib) on blood glucose levels in diabetic and nondiabetic patients in general clinical practice. J. Oncol. Pharm. Pract. 17, 197–202.

Bhargava, P. (2009). VEGF kinase inhibitors: how do they cause hypertension? Am J Physiol Regul Integr Comp Physiol 297, R1.

Blinova, K., Stohlman, J., Vicente, J., Chan, D., Johannesen, L., Hortigon-Vinagre, M.P., Zamora, V., Smith, G., Crumb, W.J., Pang, L., et al. (2017). Comprehensive Translational Assessment of Human-Induced Pluripotent Stem Cell Derived Cardiomyocytes for Evaluating Drug-Induced Arrhythmias. Toxicol. Sci. Off. J. Soc. Toxicol. 155, 234–247.

Breckenridge, R.A., Piotrowska, I., Ng, K.-E., Ragan, T.J., West, J.A., Kotecha, S., Towers, N., Bennett, M., Kienesberger, P.C., Smolenski, R.T., et al. (2013). Hypoxic Regulation of Hand1 Controls the Fetal-Neonatal Switch in Cardiac Metabolism. PLOS Biol. 11, e1001666.

Bresalier, R.S., Sandler, R.S., Quan, H., Bolognese, J.A., Oxenius, B., Horgan, K., Lines, C., Riddell, R., Morton, D., Lanas, A., et al. (2005). Cardiovascular events associated with rofecoxib in a colorectal adenoma chemoprevention trial. N. Engl. J. Med. 352, 1092–1102.

Burridge, P.W., Li, Y.F., Matsa, E., Wu, H., Ong, S., Sharma, A., Holmström, A., Chang, A.C., Coronado, M.J., Ebert, A.D., et al. (2016). Human Induced Pluripotent Stem Cell–Derived Cardiomyocytes Recapitulate the Predilection of Breast Cancer Patients to Doxorubicin–Induced Cardiotoxicity. Nat. Med. 22, 547–556.

Chen, M.H., Kerkelä, R., and Force, T. (2008). Mechanisms of Cardiac Dysfunction Associated With Tyrosine Kinase Inhibitor Cancer Therapeutics. Circulation 118, 84–95.

Cheng, H., and Force, T. (2010). Molecular Mechanisms of Cardiovascular Toxicity of Targeted Cancer Therapeutics. Circ Res 106, 21.

Chu, T.F., Rupnick, M.A., Kerkela, R., Dallabrida, S.M., Zurakowski, D., Nguyen, L., Woulfe, K., Pravda, E., Cassiola, F., Desai, J., et al. (2007). Cardiotoxicity associated with tyrosine kinase inhibitor sunitinib. Lancet 370, 2011–2019.

Davis, M.I., Hunt, J.P., Herrgard, S., Ciceri, P., Wodicka, L.M., Pallares, G., Hocker, M., Treiber, D.K., and Zarrinkar, P.P. (2011). Comprehensive analysis of kinase inhibitor selectivity. Nat Biotechnol 29, 1046–1051.

Dayon, L., and Sanchez, J.C. (2012). Relative protein quantification by MS/MS using the tandem mass tag technology. Methods Mol Biol 893, 115–127.

Doenst, T., Nguyen, T.D., and Abel, E.D. (2013). Cardiac metabolism in heart failure: implications beyond ATP production. Circ. Res. 113, 709–724.

Eid, S., Turk, S., Volkamer, A., Rippmann, F., and Fulle, S. (2017). KinMap: a web-based tool for interactive navigation through human kinome data. BMC Bioinformatics 18, 16.

Faust, P.L., and Kovacs, W.J. (2014). Cholesterol biosynthesis and ER stress in peroxisome deficiency. Biochimie 98, 75–85.

Force, T., Krause, D.S., and Van Etten, R.A. (2007). Molecular mechanisms of cardiotoxicity of tyrosine kinase inhibition. Nat Rev Cancer 7, 332–344.

Francis, J., Ahluwalia, M.S., Wetzler, M., Wang, E., Paplham, P., Smiley, S., McCarthy, P.L., Cohen, I.L., Spangenthal, E., and Battiwalla, M. (2010). Reversible cardiotoxicity with tyrosine kinase inhibitors. Clin. Adv. Hematol. Oncol. HO 8, 128–132.

Funakoshi, S., Miki, K., Takaki, T., Okubo, C., Hatani, T., Chonabayashi, K., Nishikawa, M., Takei, I., Oishi, A., Narita, M., et al. (2016). Enhanced engraftment, proliferation, and therapeutic potential in heart using optimized human iPSC-derived cardiomyocytes. Sci. Rep. 6.

Gavila, J., Seguí, M.Á., Calvo, L., López, T., Alonso, J.J., Farto, M., and Sánchez-de la Rosa, R. (2017). Evaluation and management of chemotherapy-induced cardiotoxicity in breast cancer: a Delphi study. Clin. Transl. Oncol. 19, 91–104.

Geyer, C.E., Forster, J., Lindquist, D., Chan, S., Romieu, C.G., Pienkowski, T., Jagiello-Gruszfeld, A., Crown, J., Chan, A., Kaufman, B., et al. (2006). Lapatinib plus Capecitabine for HER2-Positive Advanced Breast Cancer. N. Engl. J. Med. 355, 2733–2743.

Hamerly, G., and Elkan, C. (2004). Learning the k in k-means. In Advances in Neural Information Processing Systems 16, S. Thrun, L.K. Saul, and P.B. Schölkopf, eds. (MIT Press), pp. 281–288.

Hinrichsen, R., Hansen, A.H., Haunsø, S., and Busk, P.K. (2008). Phosphorylation of pRb by cyclin D kinase is necessary for development of cardiac hypertrophy. Cell Prolif. 41, 813–829.

Johnson, D.B., Balko, J.M., Compton, M.L., Chalkias, S., Gorham, J., Xu, Y., Hicks, M., Puzanov, I., Alexander, M.R., Bloomer, T.L., et al. (2016). Fulminant Myocarditis with Combination Immune Checkpoint Blockade. N Engl J Med 375, 1749–1755.

Karakikes, I., Ameen, M., Termglinchan, V., and Wu, J.C. (2015). Human Induced Pluripotent Stem Cell-Derived Cardiomyocytes: Insights into Molecular, Cellular, and Functional Phenotypes. Circ. Res. 117, 80–88.

Kerkela, R., Grazette, L., Yacobi, R., Iliescu, C., Patten, R., Beahm, C., Walters, B., Shevtsov, S., Pesant, S., Clubb, F.J., et al. (2006). Cardiotoxicity of the cancer therapeutic agent imatinib mesylate. Nat Med 12, 908–916.

Kerkela, R., Woulfe, K.C., Durand, J.-B., Vagnozzi, R., Kramer, D., Chu, T.F., Beahm, C., Chen, M.H., and Force, T. (2009). Sunitinib-Induced Cardiotoxicity Is Mediated by Off-Target Inhibition of AMP-Activated Protein Kinase. Clin Transl Sci 2, 15–25.

Magdy, T., Schuldt, A.J.T., Wu, J.C., Bernstein, D., and Burridge, P.W. (2018). Human Induced Pluripotent Stem Cell (hiPSC)-Derived Cells to Assess Drug Cardiotoxicity: Opportunities and Problems. Annu. Rev. Pharmacol. Toxicol. 58, null.

Mertens, A.C., Liu, Q., Neglia, J.P., Wasilewski, K., Leisenring, W., Armstrong, G.T., Robison, L.L., and Yasui, Y. (2008). Cause-specific late mortality among 5-year survivors of childhood cancer: the Childhood Cancer Survivor Study. J Natl Cancer Inst 100, 1368–1379.

Moslehi, J.J., Salem, J.-E., Sosman, J.A., Lebrun-Vignes, B., and Johnson, D.B. (2018). Increased reporting of fatal immune checkpoint inhibitor-associated myocarditis. The Lancet 391, 933.

Necela, B.M., Axenfeld, B.C., Serie, D.J., Kachergus, J.M., Perez, E.A., Thompson, E.A., and Norton, N. (2017). The antineoplastic drug, trastuzumab, dysregulates metabolism in iPSC-derived cardiomyocytes. Clin. Transl. Med. 6, 5.

Ohtani, T., Mano, T., Hikoso, S., Sakata, Y., Nishio, M., Takeda, Y., Otsu, K., Miwa, T., Masuyama, T., Hori, M., et al. (2009). Cardiac steroidogenesis and glucocorticoid in the development of cardiac hypertrophy during the progression to heart failure. J. Hypertens. 27, 1074–1083.

Orphanos, G.S., Ioannidis, G.N., and Ardavanis, A.G. (2009). Cardiotoxicity induced by tyrosine kinase inhibitors. Acta Oncol. 48, 964–970.

Robinson, E.S., Khankin, E.V., Karumanchi, S.A., and Humphreys, B.D. (2010). Hypertension induced by vascular endothelial growth factor signaling pathway inhibition: mechanisms and potential use as a biomarker. Semin. Nephrol. 30, 591–601.

Roden, D.M. (2004). Drug-induced prolongation of the QT interval. N. Engl. J. Med. 350, 1013–1022.

Rosano, G.M., Fini, M., Caminiti, G., and Barbaro, G. (2008). Cardiac metabolism in myocardial ischemia. Curr Pharm Des 14, 2551–2562.

Schmidinger, M., Zielinski, C.C., Vogl, U.M., Bojic, A., Bojic, M., Schukro, C., Ruhsam, M., Hejna, M., and Schmidinger, H. (2008). Cardiac Toxicity of Sunitinib and Sorafenib in Patients With Metastatic Renal Cell Carcinoma. J Clin Orthod 26, 5204–5212.

Shadrin, I.Y., Allen, B.W., Qian, Y., Jackman, C.P., Carlson, A.L., Juhas, M.E., and Bursac, N. (2017). Cardiopatch platform enables maturation and scale-up of human pluripotent stem cell-derived engineered heart tissues. Nat. Commun. 8, 1825.

Sharma, A., Burridge, P.W., McKeithan, W.L., Serrano, R., Shukla, P., Sayed, N., Churko, J.M., Kitani, T., Wu, H., Holmström, A., et al. (2017). High-throughput screening of tyrosine kinase inhibitor cardiotoxicity with human induced pluripotent stem cells. Sci. Transl. Med. 9.

Sogawa, K., Numayama-Tsuruta, K., Ema, M., Abe, M., Abe, H., and Fujii-Kuriyama, Y. (1998). Inhibition of hypoxia-inducible factor 1 activity by nitric oxide donors in hypoxia. Proc. Natl. Acad. Sci. U. S. A. 95, 7368–7373.

Spinelli, J.B., Yoon, H., Ringel, A.E., Jeanfavre, S., Clish, C.B., and Haigis, M.C. (2017). Metabolic recycling of ammonia via glutamate dehydrogenase supports breast cancer biomass. Science 358, 941–946.

Sun, C., Wang, L., Huang, S., Heynen, G.J.J.E., Prahallad, A., Robert, C., Haanen, J., Blank, C., Wesseling, J., Willems, S.M., et al. (2014). Reversible and adaptive resistance to BRAF(V600E) inhibition in melanoma. Nature 508, 118–122.

Uraizee, I., Cheng, S., and Moslehi, J. (2011). Reversible Cardiomyopathy Associated with Sunitinib and Sorafenib. N Engl J Med 365, 1649–1650.

Vandenberg, J.I., Perry, M.D., Perrin, M.J., Mann, S.A., Ke, Y., and Hill, A.P. (2012). hERG K(+) channels: structure, function, and clinical significance. Physiol. Rev. 92, 1393–1478.

Ventura-Clapier, R., Garnier, A., and Veksler, V. (2004). Energy metabolism in heart failure. J. Physiol. 555, 1–13.

Von Hoff, D.D., Layard, M.W., Basa, P., Davis, H.L., Jr., Von Hoff, A.L., Rozencweig, M., and Muggia, F.M. (1979). Risk factors for doxorubicin-induced congestive heart failure. Ann Intern Med 91, 710–717.

van der Vusse, G.J., Glatz, J.F., Stam, H.C., and Reneman, R.S. (1992). Fatty acid homeostasis in the normoxic and ischemic heart. Physiol. Rev. 72, 881–940.

Waxman, A.J., Clasen, S., Hwang, W.-T., Garfall, A., Vogl, D.T., Carver, J., O’Quinn, R., Cohen, A.D., Stadtmauer, E.A., Ky, B., et al. (2018). Carfilzomib-Associated Cardiovascular Adverse Events: A Systematic Review and Meta-analysis. JAMA Oncol. 4, e174519–e174519.

Will, Y., Dykens, J.A., Nadanaciva, S., Hirakawa, B., Jamieson, J., Marroquin, L.D., Hynes, J., Patyna, S., and Jessen, B.A. (2008). Effect of the multitargeted tyrosine kinase inhibitors imatinib, dasatinib, sunitinib, and sorafenib on mitochondrial function in isolated rat heart mitochondria and H9c2 cells. Toxicol. Sci. Off. J. Soc. Toxicol. 106, 153–161.

Wohlschlaeger, J., Schmitz, K.J., Takeda, A., Takeda, N., Vahlhaus, C., Stypmann, J., Schmid, C., and Baba, H.A. (2010). Reversible regulation of the retinoblastoma protein/E2F-1 pathway during “reverse cardiac remodelling” after ventricular unloading. J. Heart Lung Transplant. Off. Publ. Int. Soc. Heart Transplant. 29, 117–124.

Yuan, W., Tang, C., Zhu, W., Zhu, J., Lin, Q., Fu, Y., Deng, C., Xue, Y., Yang, M., Wu, S., et al. (2016). CDK6 mediates the effect of attenuation of miR-1 on provoking cardiomyocyte hypertrophy. Mol. Cell. Biochem. 412, 289–296.

Zhang, C., Liu, Z., Bunker, E., Ramirez, A., Lee, S., Peng, Y., Tan, A.C., Eckhardt, S.G., Chapnick, D.A., and Liu, X. (2017). Sorafenib Targets the Mitochondrial Electron Transport Chain Complexes and ATP Synthase to Activate the PINK1-Parkin Pathway and Modulate Cellular Drug Response. J. Biol. Chem. jbc.M117.783175.

Zhang, S., Liu, X., Bawa-Khalfe, T., Lu, L.-S., Lyu, Y.L., Liu, L.F., and Yeh, E.T.H. (2012). Identification of the molecular basis of doxorubicin-induced cardiotoxicity. Nat. Med. 18, 1639–1642.

Zhou, Y., Wang, L., Liu, Z., Alimohamadi, S., Yin, C., Liu, J., and Qian, L. (2017). Comparative Gene Expression Analyses Reveal Distinct Molecular Signatures between Differentially Reprogrammed Cardiomyocytes. Cell Rep. 20, 3014–3024.

