## Supplementary Figures and Text for "Adaptation of human iPSC-derived cardiomyocytes to tyrosine kinase inhibitors reduces acute cardiotoxicity via metabolic reprogramming"

[‡]Lead Contact

**Contents (Supplemental Information Inventory)**

- **Supplementary Figures Related to Figures 1-7**
  - Supplementary Figure 1A: Expression of cardiomyocyte markers and sarcomere structure in Cor.4U hiPSC-CMs (with Figure 1).
  - Supplementary Figure 1B: Contractility response of human iPSC-derived cardiomyocytes to four TKIs (with Figure 1).
  - Supplementary Figure 1C: Experimental design of phenotypic, transcriptomic and proteomic studies (with Figure 1 and STAR method)
  - Supplementary Figure 2A: Correlation between replicates of RNAseq data (with Figure 2).
  - Supplementary Figure 2B: Quantification of gene expression changes in the vehicle-only control and drug treatment conditions (with Figure 2).
  - Supplementary Figure 2C: Fraction of proliferating cells in hiPSC-CMs over time (with Figure 2).
  - Supplementary Figure 2D: Relation of time-dependent and dose-dependent gene expression in response to Sunitinib treatment (with Figure 2).
  - Supplementary Figure 2E: Correlation of differential gene expression between RNAseq and qRT-PCR measurements (with Figure 2).
  - Supplementary Figure 2F: Drug treatment and sample collection schedules for RNA-seq and metabolite studies on conditioned media (with STAR method).
  - Supplementary Figure 3: Principal component analysis of TKI- and Doxorubicin-induced RNA differential expression (with Figure 3)
  - Supplementary Figure 4A: Design of proteomics studies (with Figure 4 and STAR method).
  - Supplementary Figure 4B: Three views of repeatability of proteomics (with Figure 4).
  - Supplementary Figure 4C: Number of differentially expressed proteins in proteomics experiments (with Figure 4).
  - Supplementary Figure 4D: Determining k for k-means clustering (with Figure 4).
  - Supplementary Figure 5A: Principal component analysis of RNA and protein differential expression (with Figure 5).
  - Supplementary Figure 5B: Comparison of differential expression in RNA and protein in response to TKI treatments (with Figure 5).
  - Supplementary Figure 5C: Expression changes in enzymes in the cholesterol synthesis pathway following Lapatinib treatment (with Figure 5).
  - Supplementary Figure 5D: Expression changes in sarcomeric components following Sorafenib treatment (with Figure 5).
  - Supplementary Figure 6: Effect of Sorafenib on mitochondrial respiration in a line of male hiPSC-CMs (with Figure 6).
  - Supplementary Figure 7: Changes in normalized ATP levels in response to 2DG alone (with Figure 7).
- **Supplementary Tables 1-8:**
  - Supplementary Table 1: Cardiotoxic clinical phenotypes induced by four TKIs (with Figure 1)
  - Supplementary Table 2: Pharmacological properties of four TKIs (with Figure 1)

- Supplementary Table 3: Genes in clusters with different enriched GO terms
- Supplementary Table 4: Genes for qRT-PCR validation (with Supplementary Figure 5)
- Supplementary Table 5: Proteins in clusters with different enriched GO terms
- Supplementary Table 6: Kinases expressed by Cor.4U hiPSC-CMs
- Supplementary Table 7: Detection of TKI target mRNA or protein in Cor.4U hiPSC-CM cells
- Supplementary Table 8: Metabolome of Cor.4U hiPSC-CMs following treatment with Sorafenib

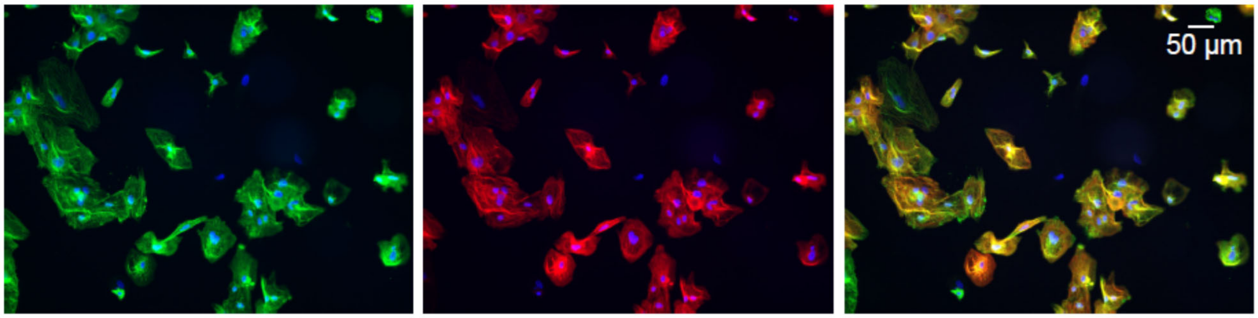


**Supplementary Figure 1A: Expression of cardiomyocyte markers and sarcomere structure in Cor.4U hiPSC-CM cells (with Figure 1).** Cells were stained with α-actinin (green), Troponin T2 (red) and DAPI for nuclei (blue) and imaged. These are lower magnification images of the same cells shown in Figure 1A.

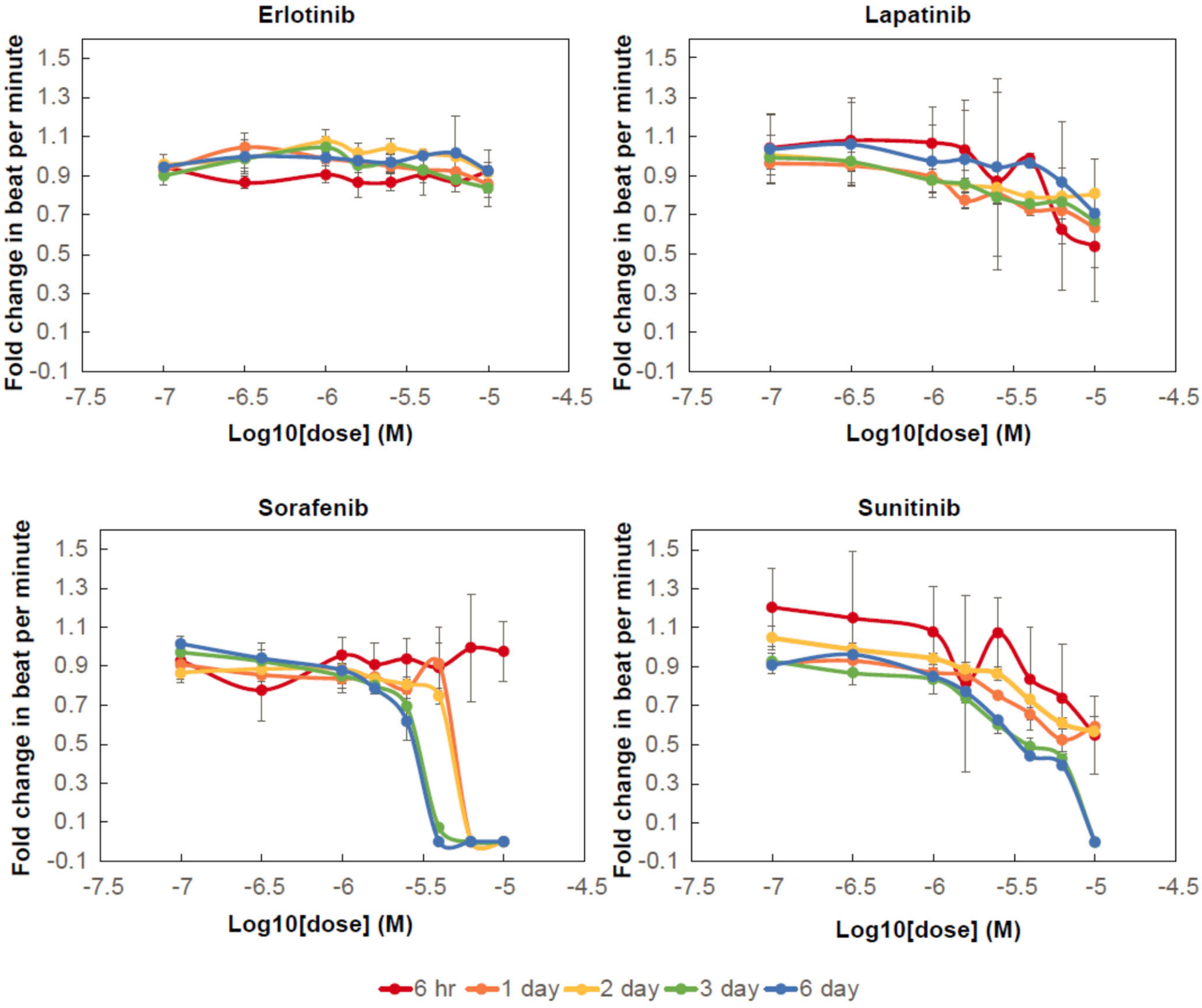


**Supplementary Figure 1B: Contractility response of human iPSC-derived cardiomyocytes to four TKIs (with Figure 1).** Cells were stained with a calcium dye using the EarlyTox kit and beat frequency was quantified based on the oscillation frequency of calcium signaling. Fold change in beat per minute was normalized to that of vehicle control and quantified for all drug treatment.

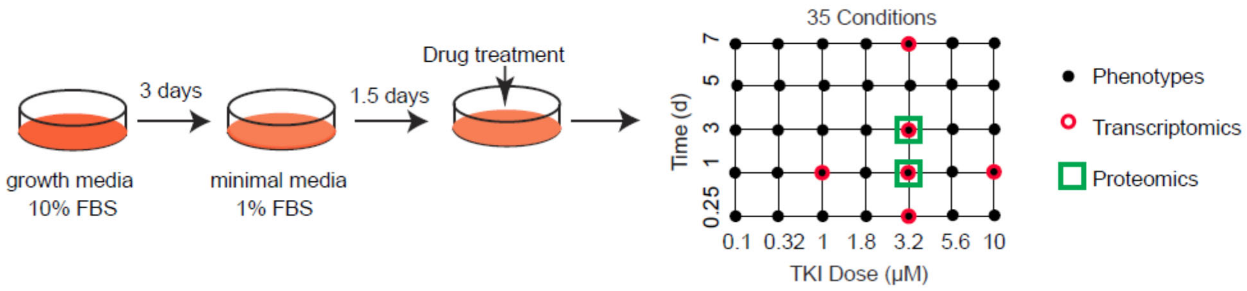


**Supplementary Figure 1C: Experimental design of phenotypic, transcriptomic and proteomic studies (with Figure 1 and STAR method).** Cells were cultured and treated as shown, left. The right panel shows the drug doses and times used for analysis of cellular phenotypes (35 conditions per drug assayed for mitochondrial membrane potentials and ATP levels), RNA-seq (6 conditions per drug) or proteomics (2 conditions per drug).

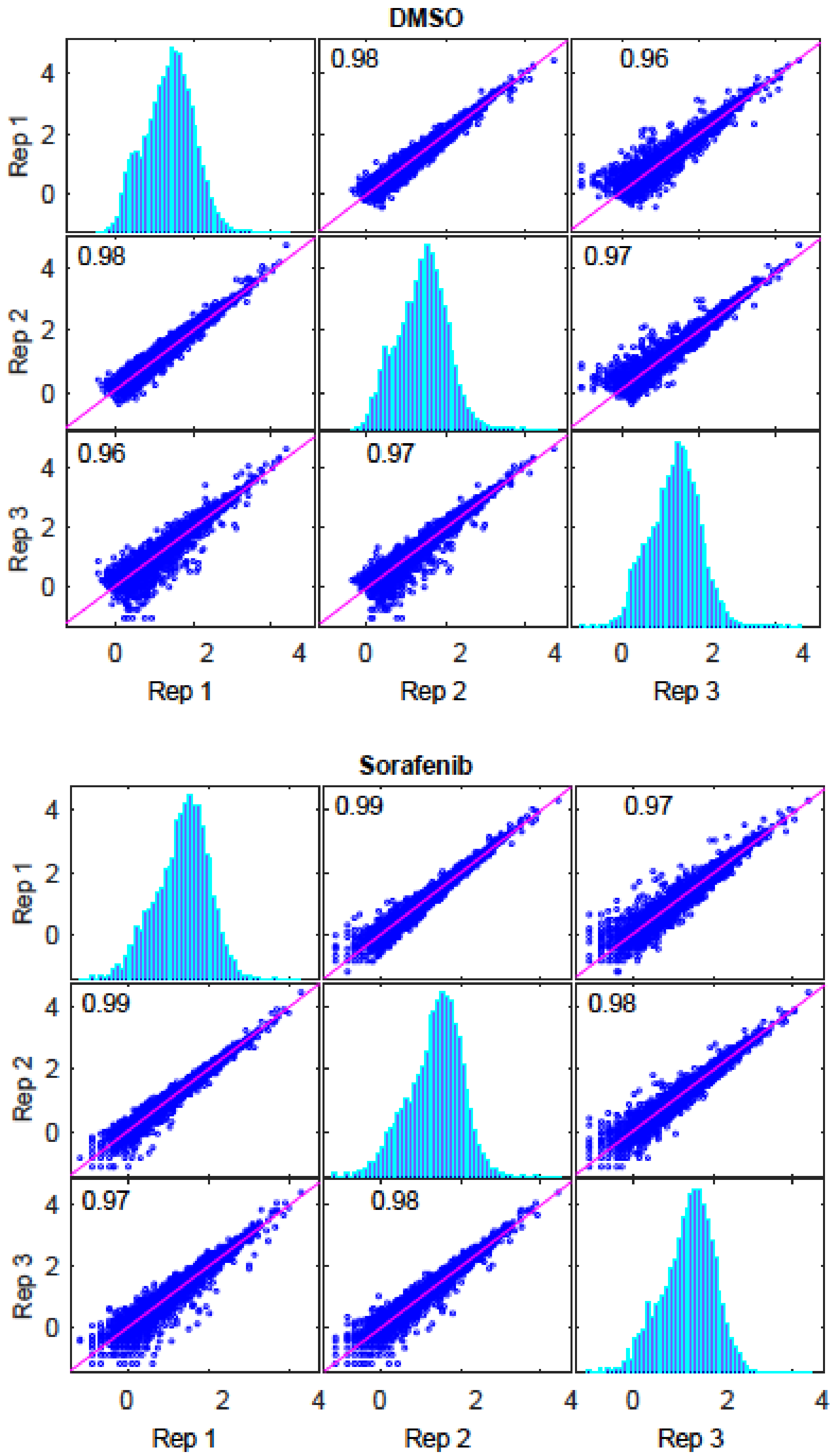


**Supplementary Figure 2A: Correlation between replicates of RNAseq data (with Figure 2).** Pearson correlation of TMM-normalized counts data between replicates of DMSO treatment at 24 hours (top) and Sorafenib treatment at 10μM and 24 hours (bottom).

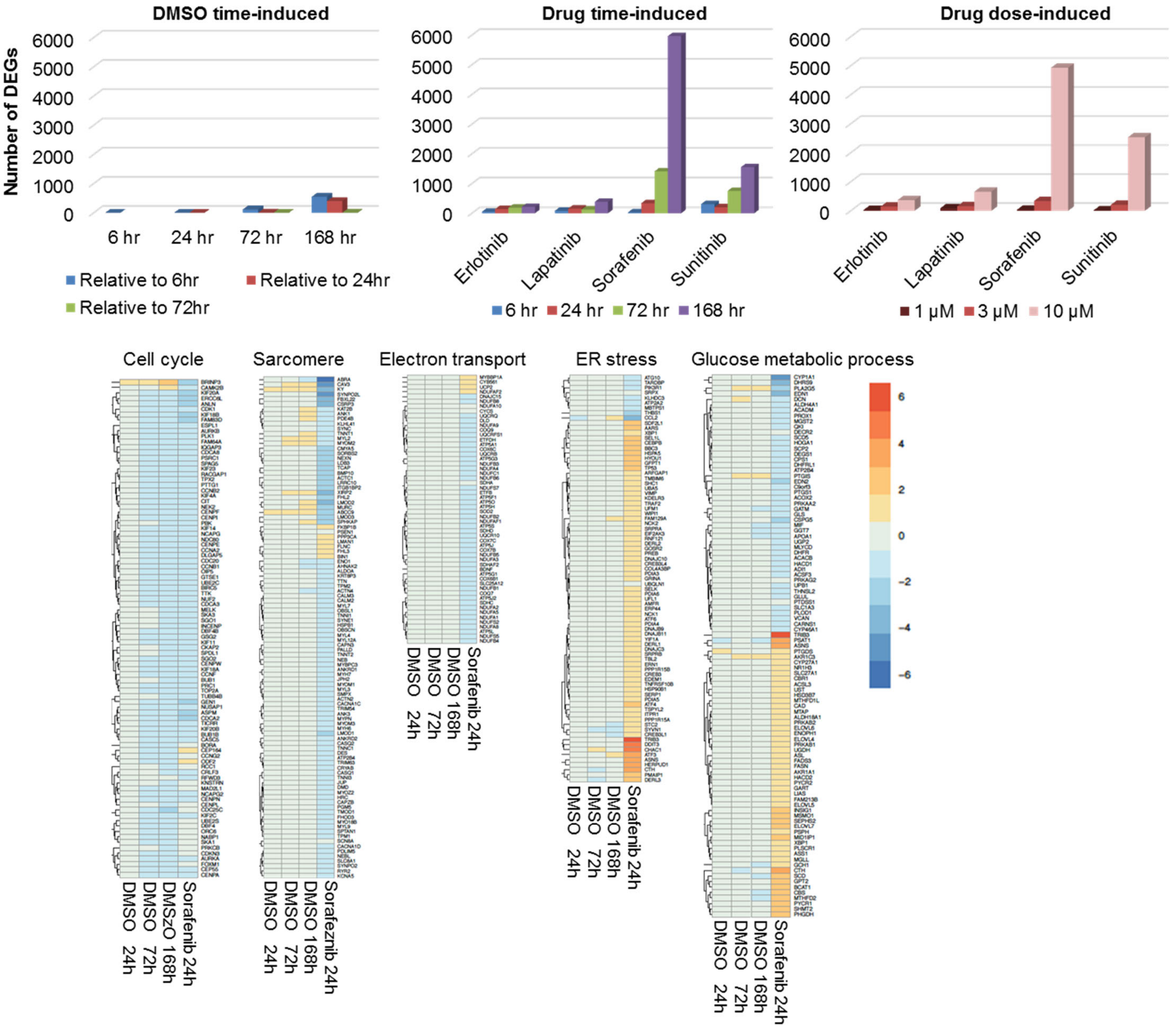


**Supplementary Figure 2B: Quantification of gene expression changes in the vehicle-only control and drug treatment conditions (with Figure 2).** The bar graphs represent the number of differentially expressed genes (DEGs) in the DMSO vehicle-only control at each time point, and in each drug treatment condition relative to vehicle-only controls were plotted. Below, the Gene Ontology (GO)-term enrichment analysis was performed based on the DEGs. Log2-based fold-changes of genes in selected enriched GO terms was plotted. Fold-changes in the DMSO conditions were relative to DMSO at 6 hours.

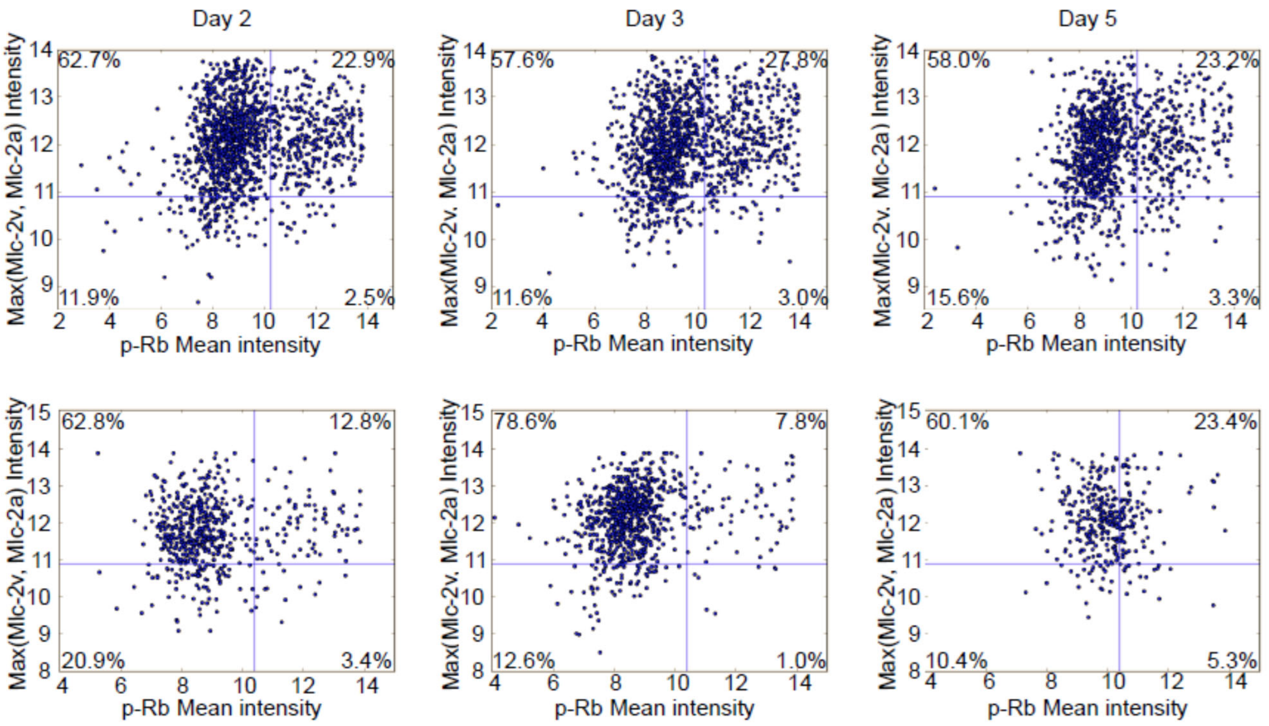


**Supplementary Figure 2C: Fraction of proliferating cells in hiPSC-CM culture over time (with Figure 2).** hiPSC-CM cells were co-stained with cardiomyocyte markers (Mlc-2v and Mlc-2a) and a cell proliferation marker (pRb). Logarithmic transformed fluorescent intensity for either marker was plotted. The y-axis corresponds to the maximal intensity of either Mlc-2v (ventricular cardiomyocyte marker) or Mlc-2a (atrial cardiomyocyte marker). The gate for mlc-max+ cells is based on positive staining for either Mlc-2v or Mlc-2a. Top row: DMSO treated, Bottom row: Sorafenib treated at 3 μM for 2, 3 and 5 days.

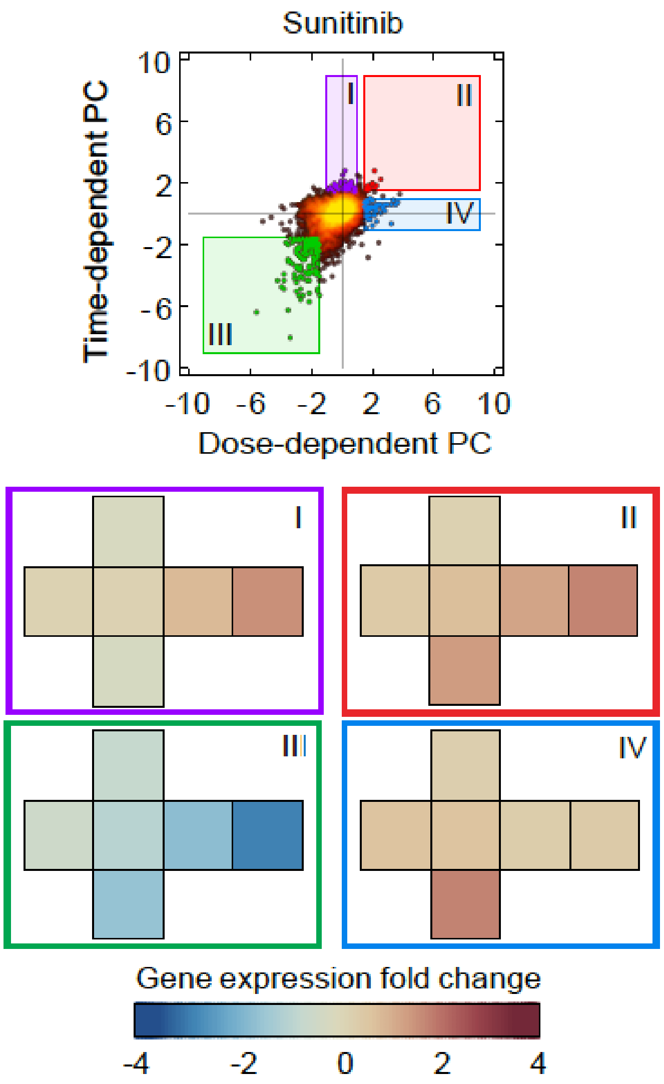


**Supplementary Figure 2D: Relation of time-dependent and dose-dependent gene expression in response to Sunitinib treatment (with Figure 2).** Genes in four different regions of the Sunitinib principal component plot (same as in Figure 2D) were selected based on their locations in PC-space. Below, the average expression changes in each region were plotted in heat maps following the cross-like experimental design.

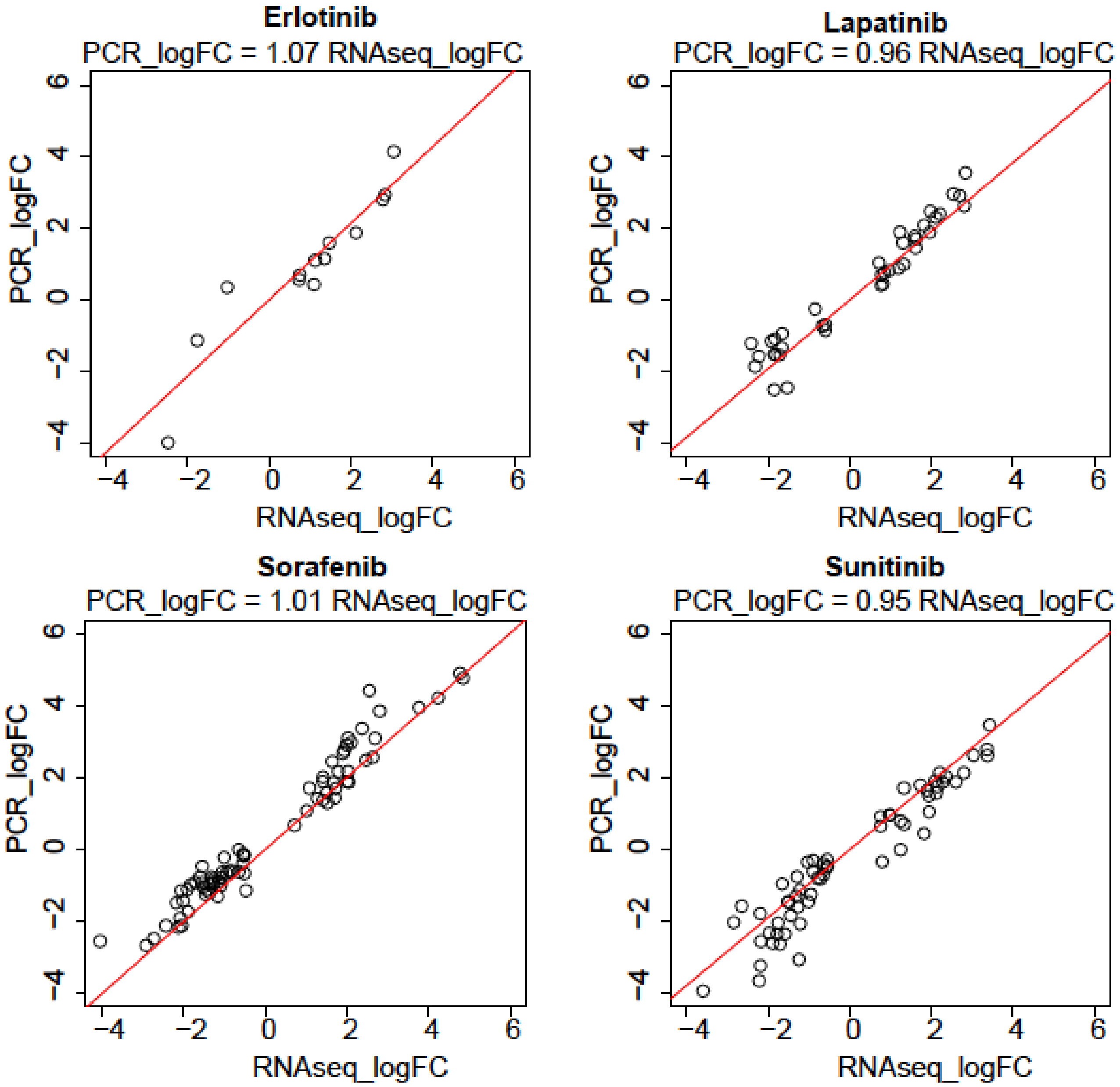


**Supplementary Figure 2E: Correlation of differential gene expression between RNAseq and qRT-PCR measurements (with Figure 2).** $Log_2$-based fold-changes measured in RNAseq and qRT-PCR under drug treatment of 10 μM and 24 hours were plotted in black circles. Data were fit by linear regression (red line) and the equation of each fit is above each individual plot.

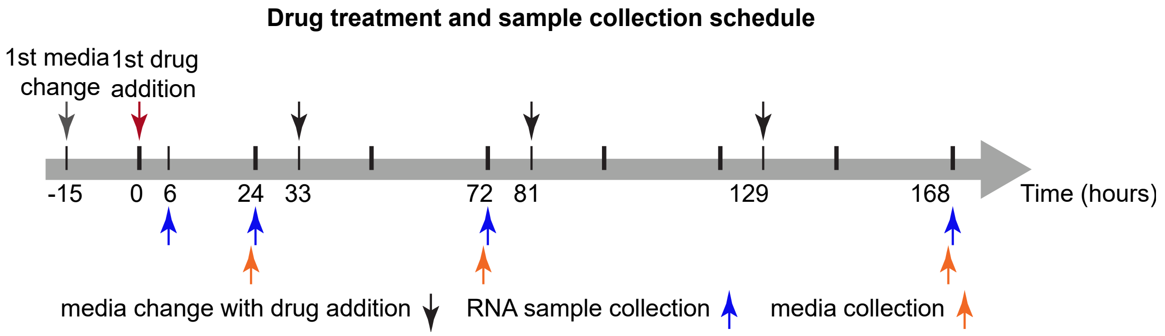


**Supplementary Figure 2F: Drug treatment and sample collection schedules for RNA-seq and metabolite studies on conditioned media (with Figure 2 and STAR method)**

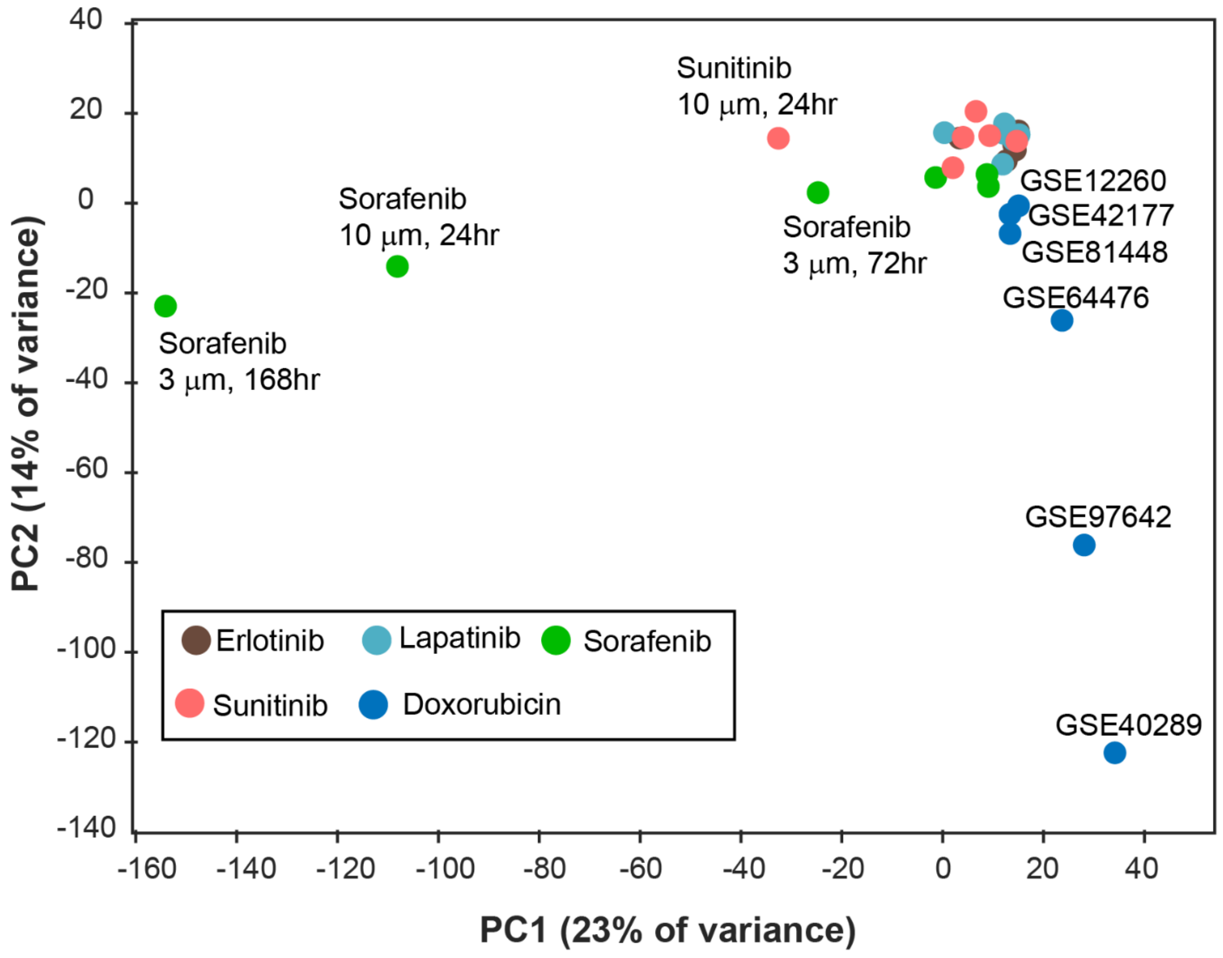


**Supplementary Figure 3: Principal component analysis of TKI- and Doxorubicin-induced RNA differential expression (with Figure 3)**. PCA was performed for $log_2$-based fold-changes measured by RNAseq for TKI treatments and for Doxorubicin treatments mined from the GEO database (with the dataset labelled as "GSE…"). Data were plotted against the first two PCs, which explained 23.3% and 14.2% of variance, respectively.

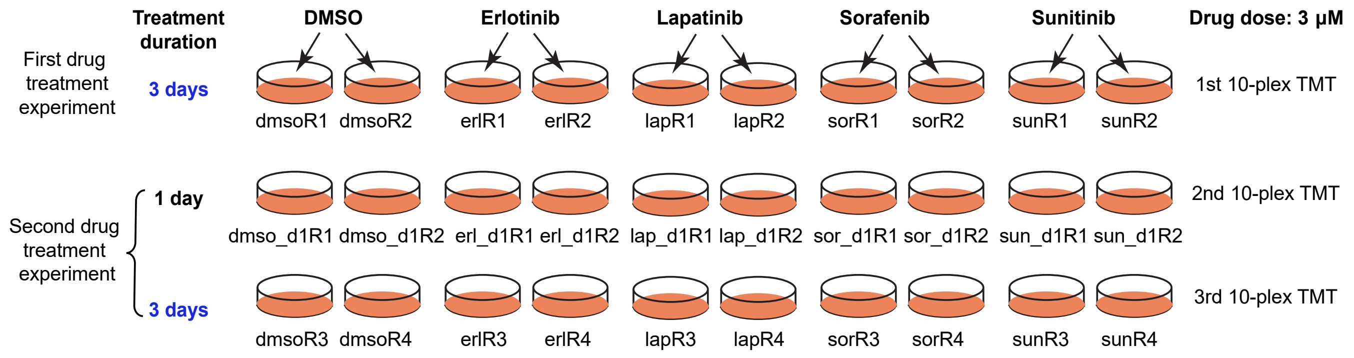


**Supplementary Figure 4A: Design of proteomics studies (with Figure 4 and STAR method).** The text at the bottom of each plate shows the code used to identify the different samples in primary data files.

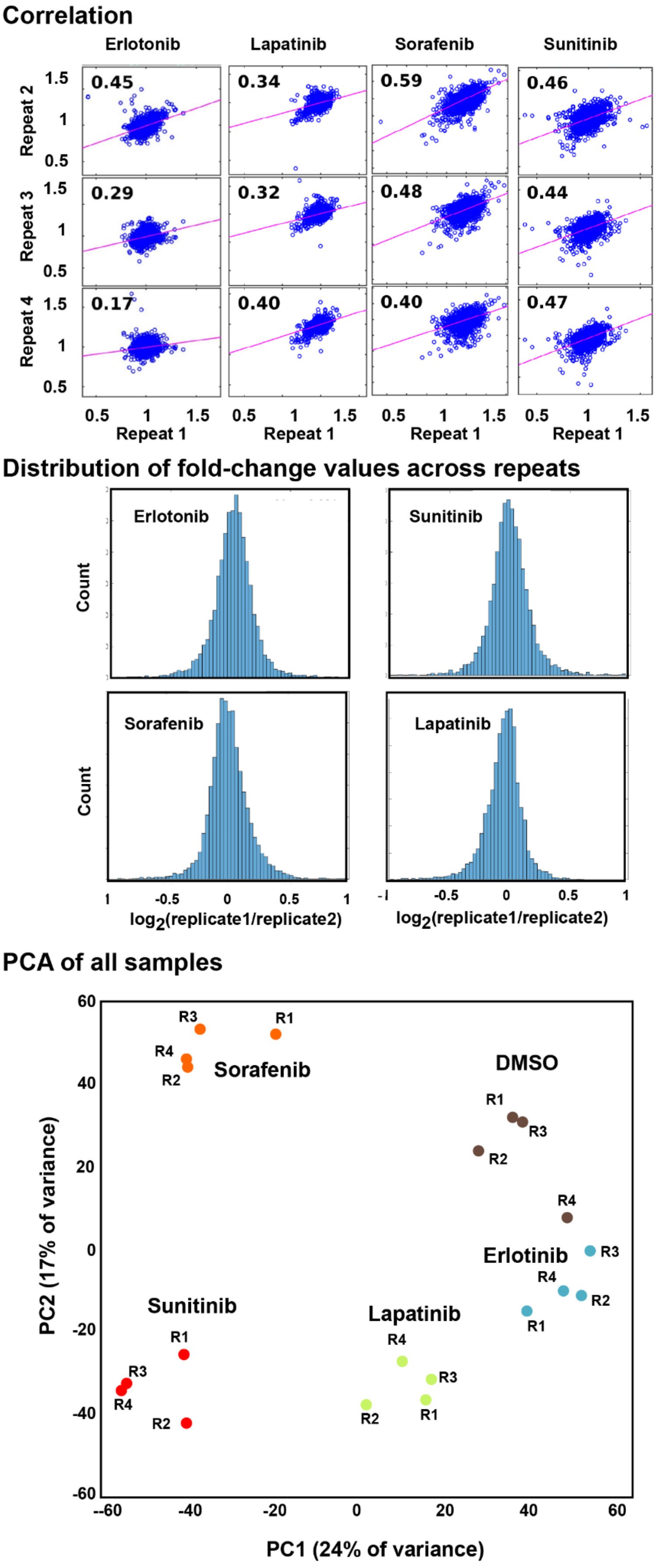


**Supplementary Figure 4B: Three views of repeatability of proteomics (with Figure 4).**

Top Panel: Pearson correlation between biological repeats of drugs treated at 3 μM and 72 hours.

Middle Panel: Fold change difference between biological replicates in proteomics experiments. Histograms of the fold change of normalized protein levels between each biological replicate are shown as a measure of reproducibility. Perfect data would have a ratio of 1 (or $\log_2 1 = 0$). Data were obtained from cells treated with the drugs indicated at 3 μM for 72 hours.

Bottom Panel: PCA of all proteomics measurements demonstrating good clustering of repeats.

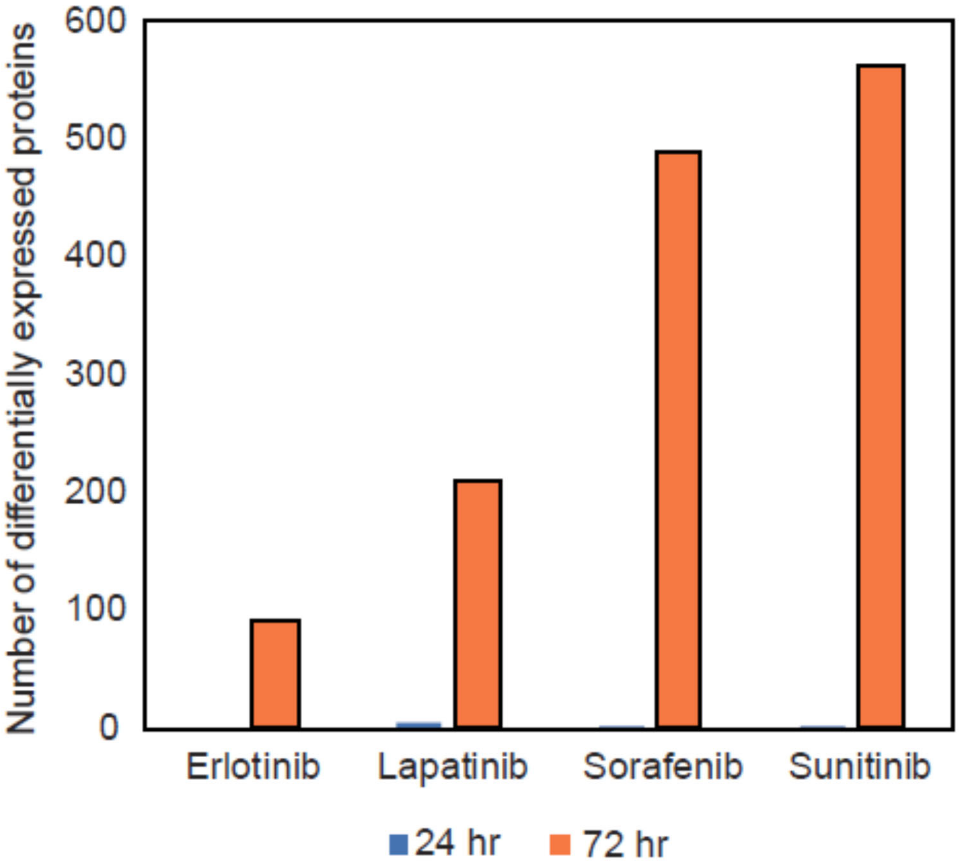


**Supplementary Figure 4C: Number of differentially expressed proteins in proteomics experiments (with Figure 4).** Based on protein-wise ANOVA statistical test with multiple test correction, number of differentially expressed proteins (FDR ≤ 0.05, fold change ≥ 1.2) was plotted.

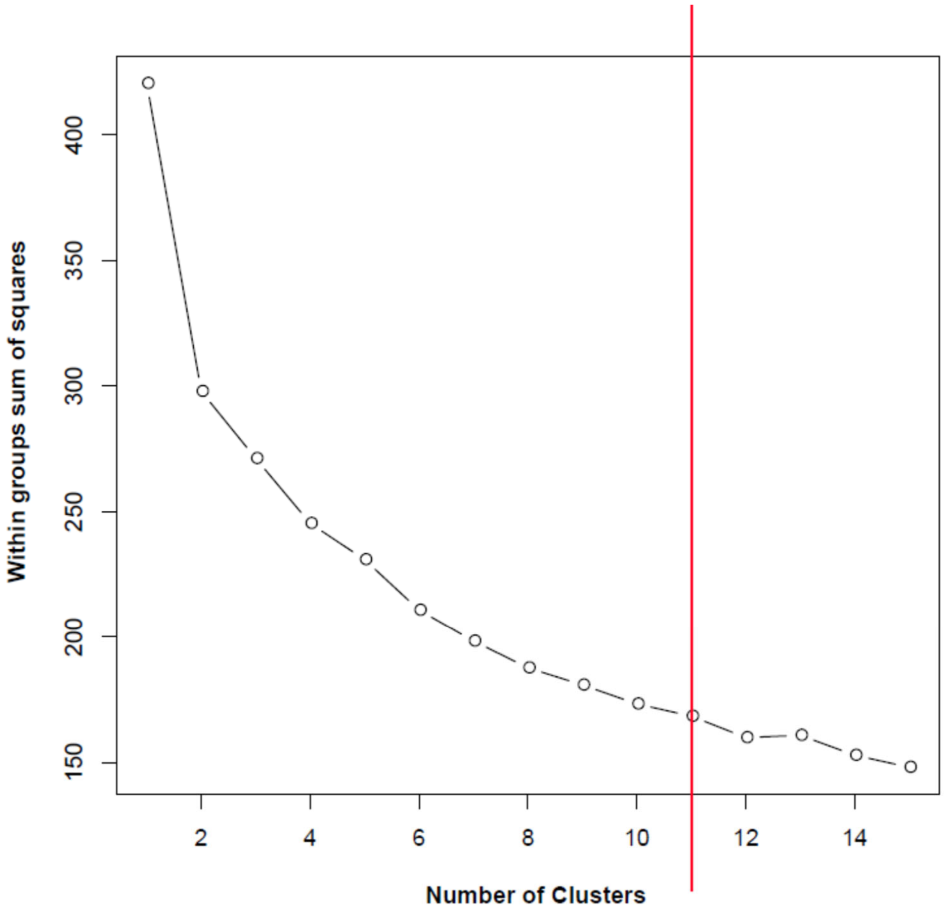


**Supplementary Figure 4D: Determining k for k-means clustering (with Figure 4).** Within-group sum of squares was calculated using k from 2-15 for the proteomics data. k=11 was selected as further increasing k did not significantly decrease within-group sum of squares.

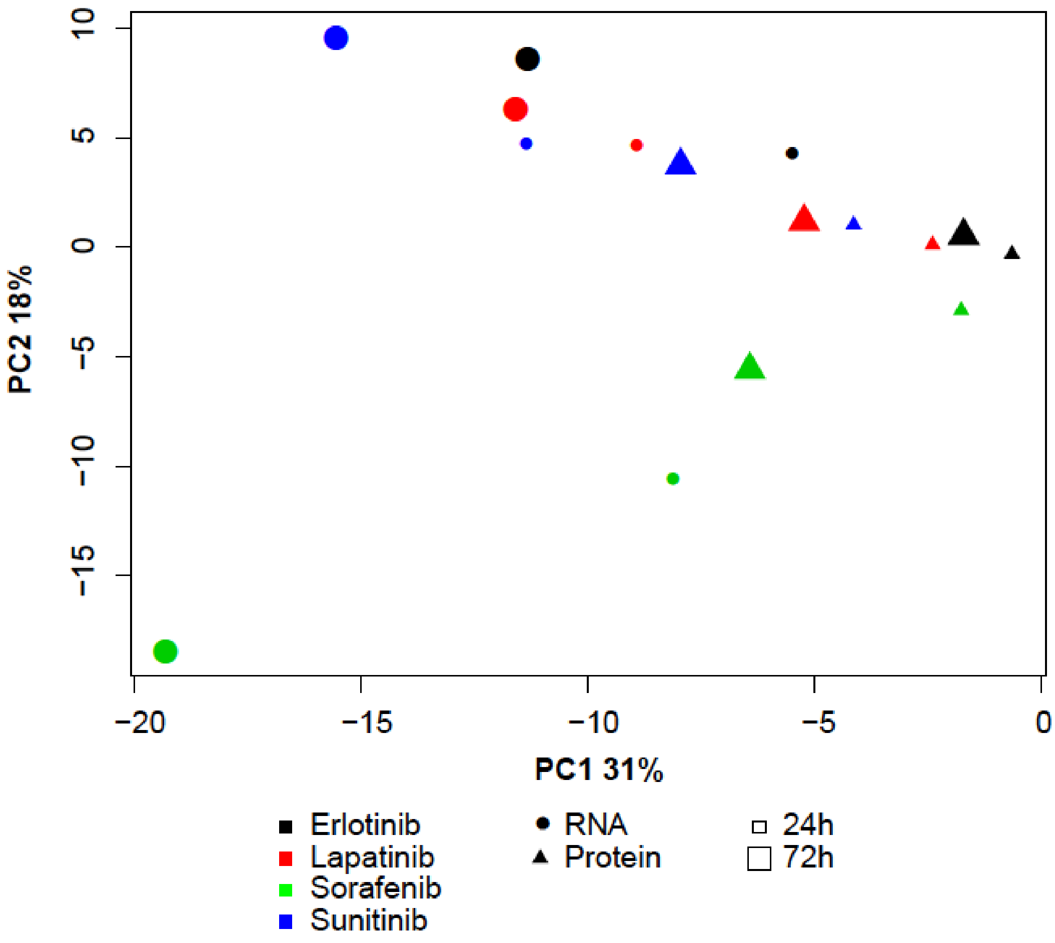


**Supplementary Figure 5A: Principal component analysis of RNA and protein differential expression (with Figure 5).** PCA was performed for $log_2$-based fold changes of both RNA and protein levels following drug treatment at one dose (3 μM) and two time points (24hr and 72hr). Data were plotted against the first two PCs, which explained 31% and 18% of the variance respectively.

| | number of DEG | number of DEP | number of overlap between DEG and DEP |
|---|---|---|---|
| Erl_3_72 | 189 | 90 | 3 |
| Lap_3_72 | 127 | 209 | 15 |
| Sor_3_72 | 1412 | 490 | 98 |
| Sun_3_72 | 753 | 562 | 18 |

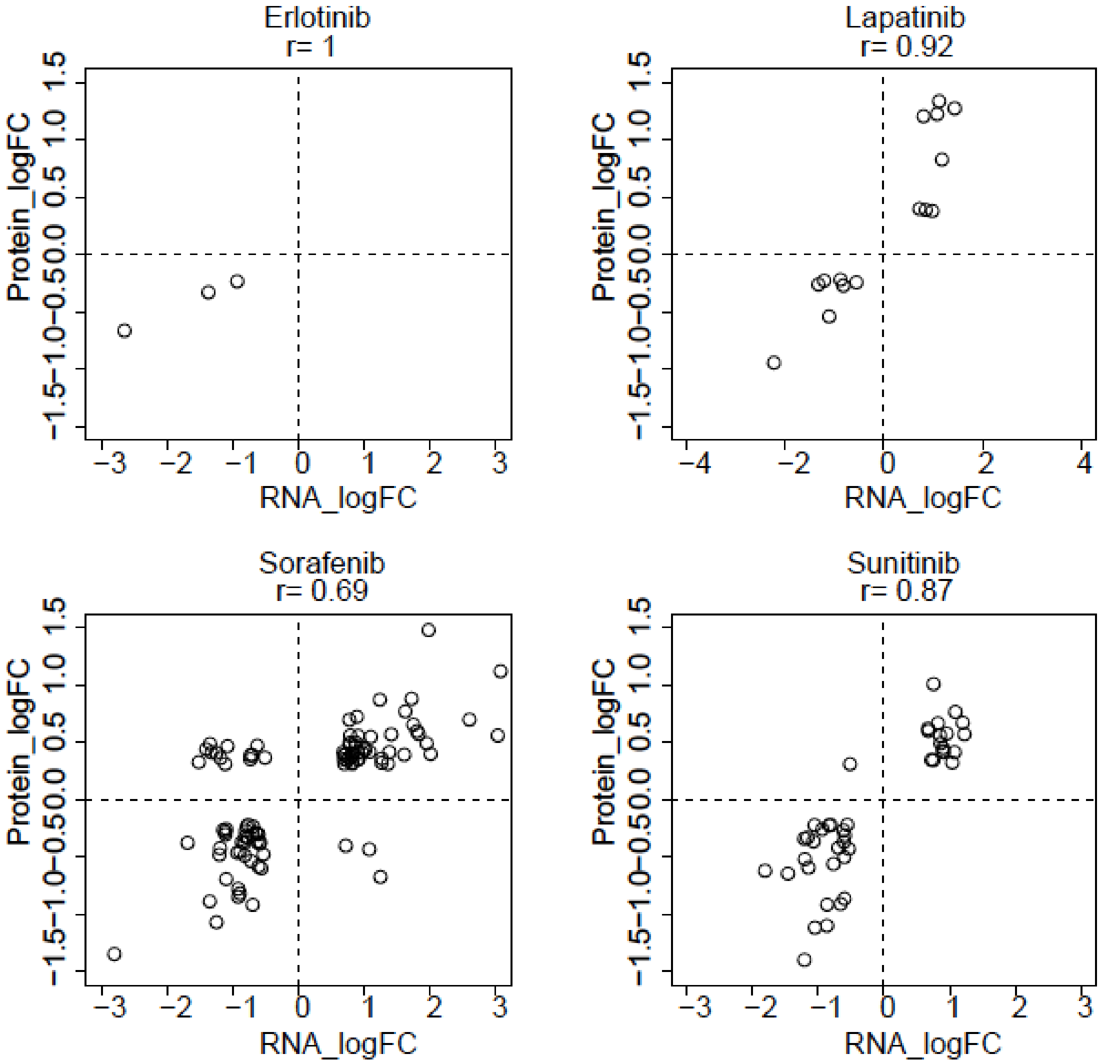


**Supplementary Figure 5B: Comparison of differential expression in RNA and protein in response to TKI treatments (with Figure 5).** The table displays the intersection of significantly differentially expressed genes (DEG) in RNAseq and differentially expressed proteins (DEP) in mass spectrometry experiments induced by TKIs. Lower four panels display the overlapping genes between DEG and DEP, $log_2$-based fold-changes from the two omics experiments were plotted. Pearson correlation coefficient (r) for each drug is reported.

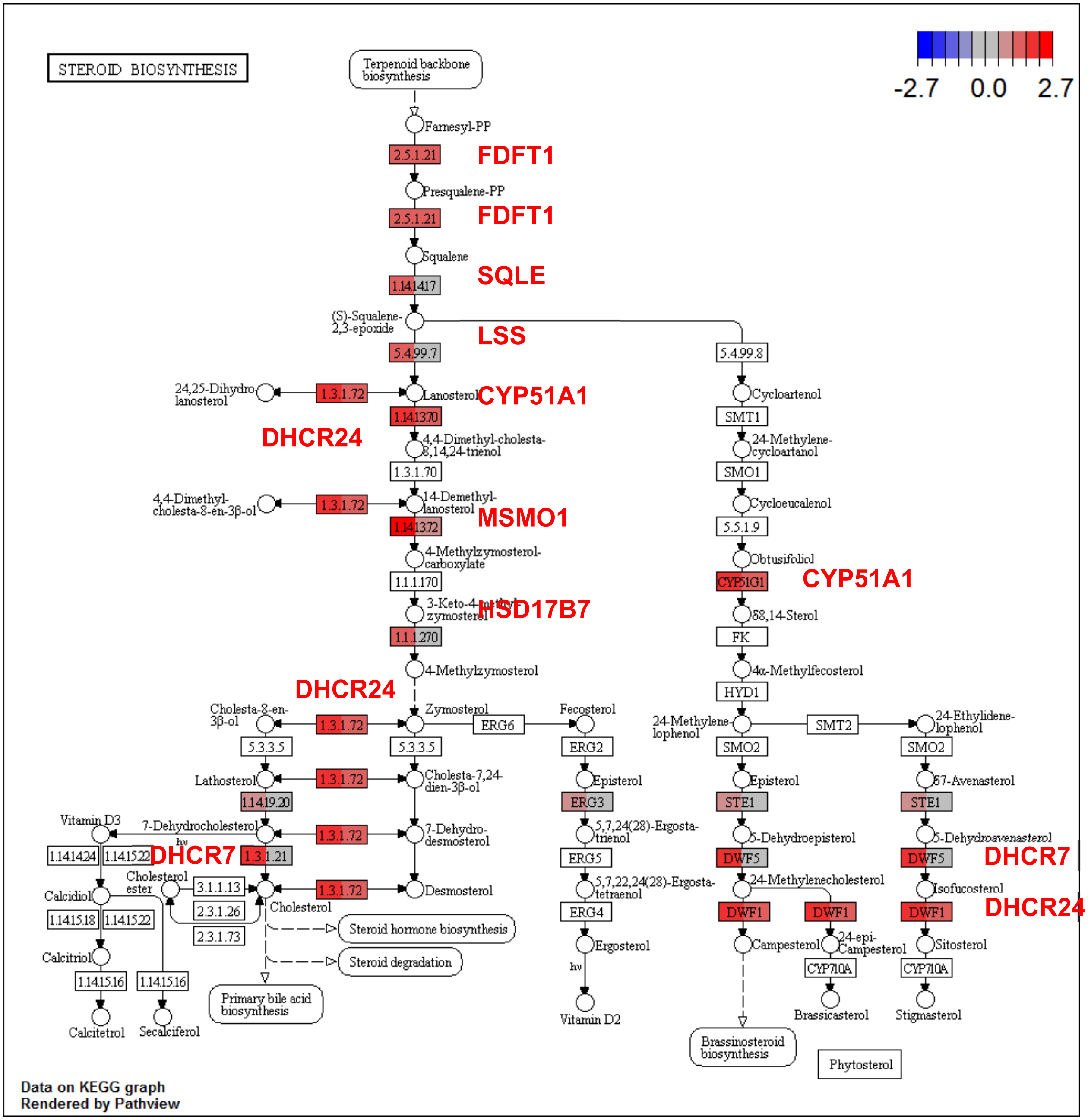


**Supplementary Figure 5C: Expression changes in enzymes in the cholesterol synthesis pathway following Lapatinib treatment (with Figure 5).** Log2-based fold-changes for RNA and protein were mapped onto the KEGG pathway for steroid biosynthesis. Red text denotes genes for which differential expression data are shown. For each gene, the left half of the box was color-coded to encode fold-changes in gene expression when treated at with 10 μM Lapatinib for 24 hr and the right half fold-changes in protein abundance when treated with 3 μM Lapatinib for 72 hr.

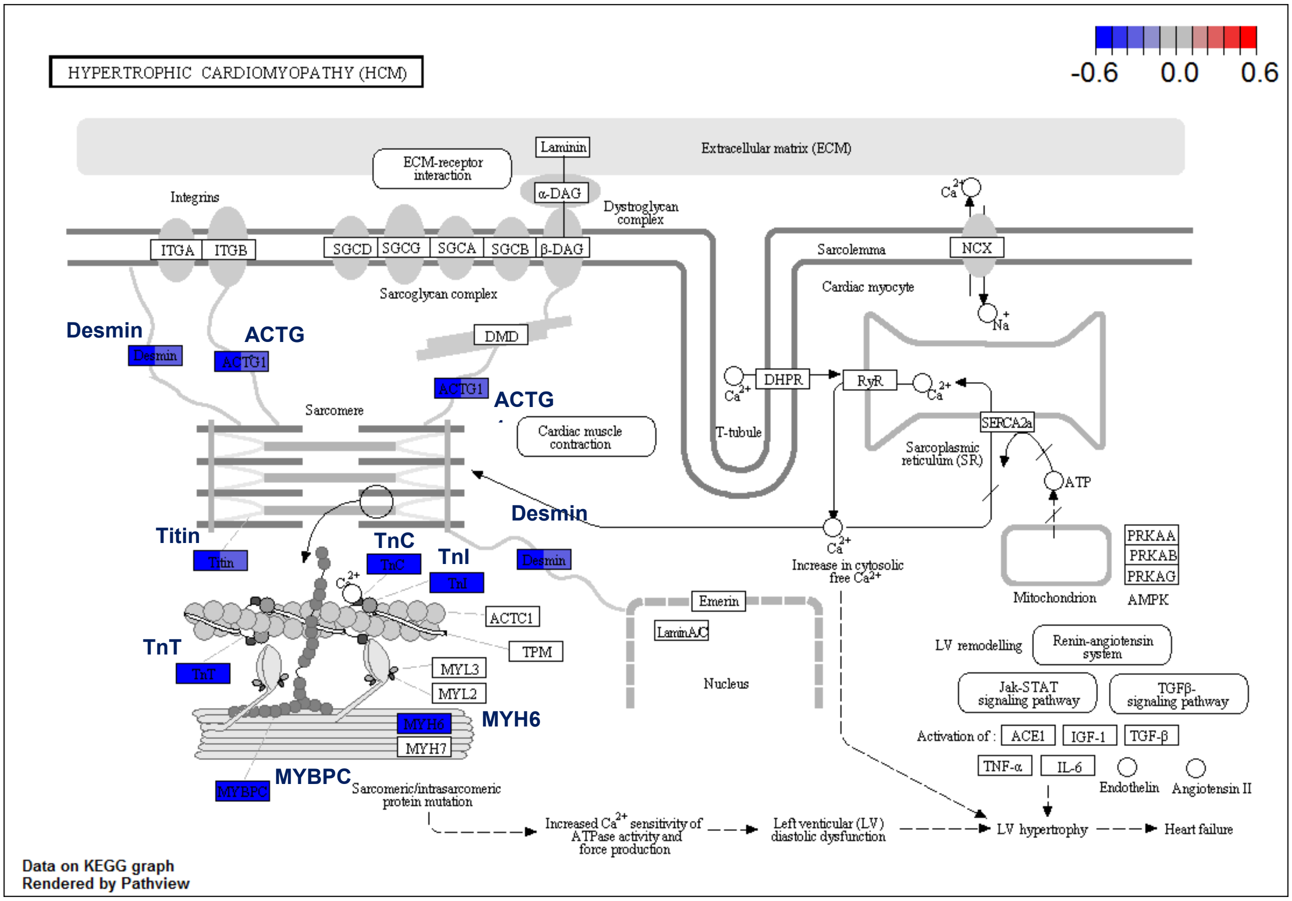


**Supplementary Figure 5D: Expression changes in sarcomeric components following Sorafenib treatment (with Figure 5).** $Log_2$-based fold-changes for RNA and protein were mapped onto the KEGG pathway for hypertrophic cardiomyopathy. For each colored box, the left half was color-coded for fold-changes in gene expression when treated at with 10 μM Sorafenib for 24 hr and the right half for fold-changes in protein abundance when treated with 3 μM Sorafenib for 72 hr.

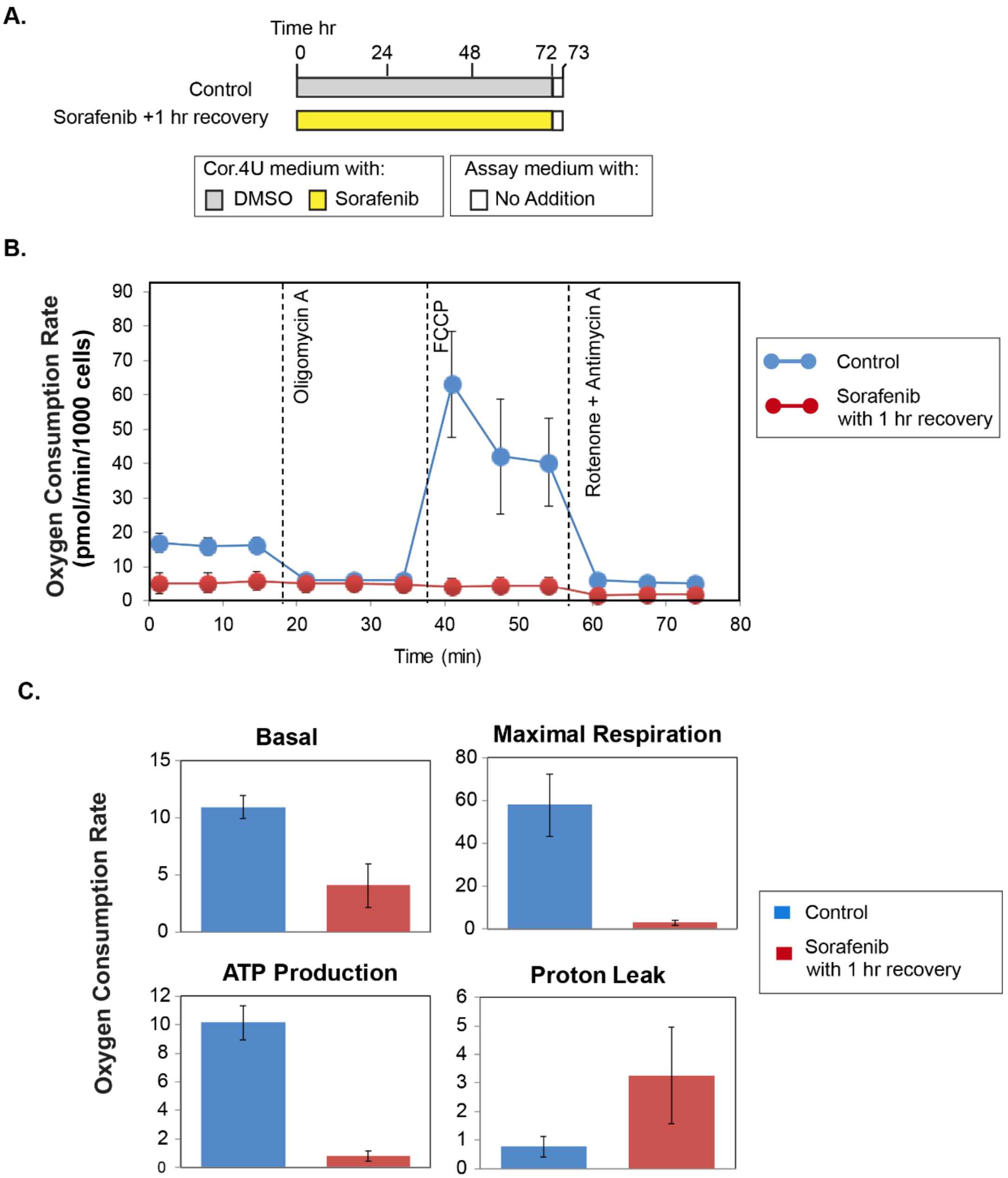


**Supplementary Figure 6: Effect of Sorafenib on mitochondrial respiration in a line of male hiPSC-CMs (with Figure 6). A.** Experimental design for testing the effect of Sorafenib on mitochondrial respiration in a line of male hiPSC-CMs is shown. **B.** The oxygen consumption rate (OCR) in male hiPSC-CMs was measured at baseline and following successive addition of Oligomycin A, FCCP and Rotenone plus Antimycin A (see text for details). Conditions correspond to those described in the design above. This experiment was repeated twice with 6 replicates per condition in each experiment; data were representative of one experiment. Error bars are standard deviation. **C.** Metabolic parameters derived from analysis of the OCR data shown as bar graphs below. Error bars are standard deviation.

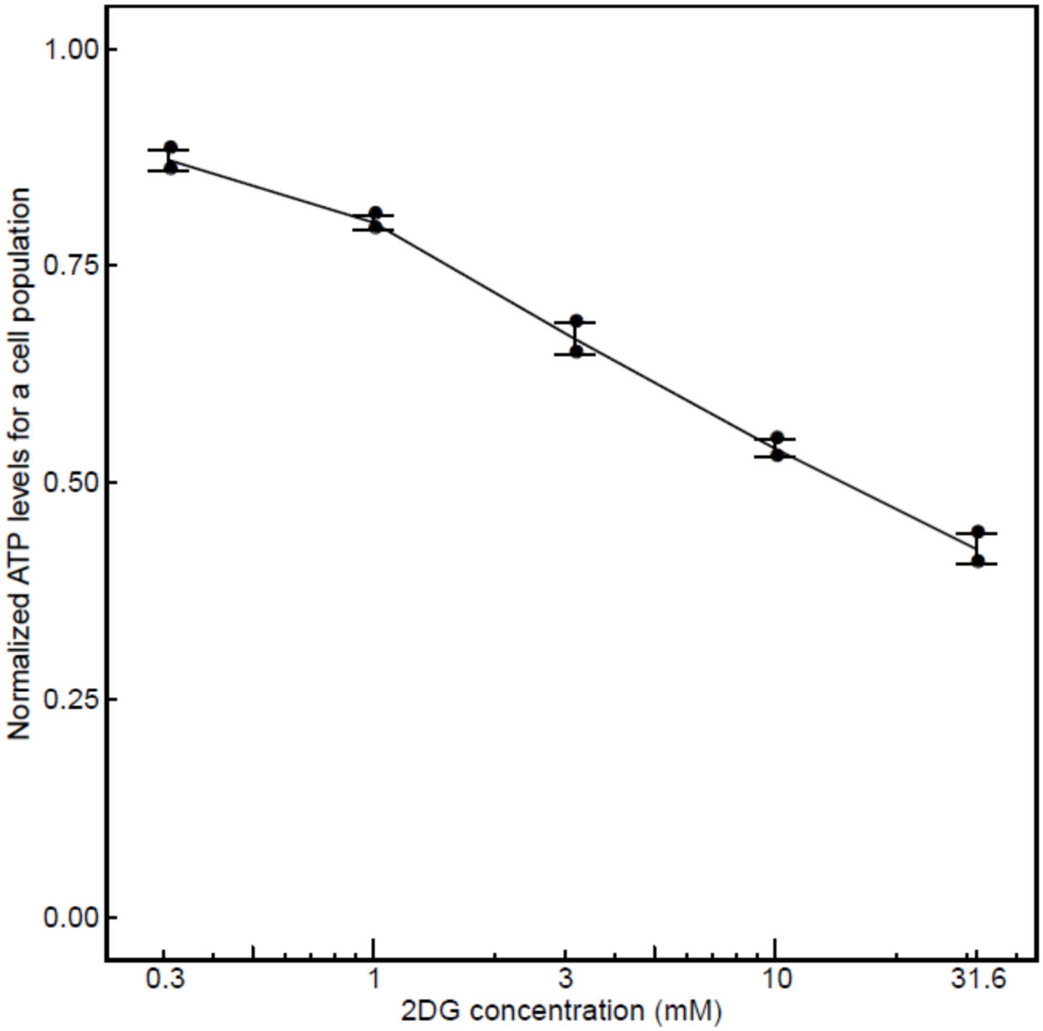


**Supplementary Figure 7: Changes in normalized ATP levels in response to 2DG alone (with Figure 7).** ATP levels of a cell population treated with 2DG at different doses were normalized to that of a vehicle-only treated control. Changes in normalized ATP levels of a cell population were plotted against 2D concentration.
