## Supplementary material for "Adaptation of human iPSC-derived cardiomyocytes to tyrosine kinase inhibitors reduces acute cardiotoxicity via metabolic reprogramming": Key Resources Table

| REAGENT or RESOURCE | SOURCE | IDENTIFIER |
|---|---|---|
| **Antibodies** | | |
| Mouse monoclonal anti-Cardiac Troponin T | Thermofisher | Cat#MA5-12960; RRID: AB-11000742; Clone: 13-11; Lot: QC1998316A |
| Mouse monoclonal anti-Sarcomeric Alpha Actinin | Abcam | Cat#ab9465; RRID: AB-307264; Clone: EA-53 |
| Rabbit monoclonal anti-Cardiac Troponin T | GeneTex | Cat#GTX62263; RRID: AB-10620847; Clone: EPR3696 |
| Rabbit monoclonal anti-Phospho-Rb (Ser807/811) with Alexa Fluor® 555 Conjugate | Cell signaling technology | Cat#8957; Clone: D20B12 |
| Rabbit polyclonal anti-PEX14 | GeneTex | Cat# GTX129230 |
| Mouse monoclonal anti-Mlc-2a with ATTO® 647N | Synaptic systems | Cat# 311011AT1 |
| Rabbit polyclonal anti-Mlc-2v | Proteintech | Cat# 10906-1-AP |
| **Bacterial and Virus Strains** | | |
| **Biological Samples** | | |
| **Chemicals, Peptides, and Recombinant Proteins** | | |
| 2-deoxy-D-glucose | Sigma-Aldrich | Cat#D8375; CAS: 154-17-6 |
| Antimycin A | Sigma-Aldrich | Cat#A8674; CAS: 1397-94-0 |
| Calcein, AM | Thermo Fisher Scientific | Cat#C3099 |
| Carbonyl cyanide 4-(trifluoromethoxy) phenylhydrazone, or FCCP | Sigma-Aldrich | Cat#C2920; CAS: 370-86-5 |
| Erlotinib | Haoyuan Chemexpress | Cat#HY-50896; CAS: 183321-74-6 |
| Hoeschst 33342, Trihydrochloride, Trihydrate | Thermo Fisher Scientific | Cat#H3570 |
| Lapatinib | Haoyuan Chemexpress | Cat#HY-50898; CAS: 231277-92-2 |
| MitoTracker Deep Red | Thermo Fisher Scientific | Cat#M22426 |
| Oligomycin A | Sigma-Aldrich | Cat#75351; CAS: 579-13-5 |
| Rotenone | Sigma-Aldrich | Cat#R8875; CAS: 83-79-4 |
| Sorafenib | LC Laboritories | Cat#S-8599; CAS: 284461-73-0 |
| Sunitinib | Haoyuan Chemexpress | Cat#HY-10255A; CAS: 557795-19-4 |
| Tetramethylrhodamine ethyl ester perchlorate | Sigma-Aldrich | Cat#87917; CAS:115532-52-0 |
| **Critical Commercial Assays** | | |
| CellTiter-Glo® Luminescent Cell Viability Assay | Promega | Cat#G7573 |

| | | |
|---|---|---|
| Custom RT2 profiler PCR arrays | Qiagen | Cat#CAPH13532 |
| Odyssey blocking buffer | Li-Cor | Cat#927-40000 |
| RT2 First Strand Kit | Qiagen | Cat#330404 |
| RT² SYBR Green ROX qPCR Mastermix | Qiagen | Cat#330524 |
| TMT10plex™ Isobaric Label Reagent Set | Thermo Fisher Scientific | Cat#90406 |
| TruSeq® Stranded Total RNA Library Prep Gold | Illumina | Cat#20020599 |
| **Deposited Data** | | |
| Pre-processed proteomics data | This paper | Synapse: syn7079983 |
| Quantitative real-time PCR data | This paper | Synapse: syn7079983 |
| Raw and analyzed RNAseq data | This paper | GEO: GSE114686 (Reviewer token: evmjucskrbobfwp) Synapse: syn7079983 |
| **Experimental Models: Cell Lines** | | |
| Cor.4U cells (female) | Ncardia (previously Axiogenesis) | Cat# Ax-B-HC02-4M (from Axiogenesis) RRID:CVCL_Y550 |
| Cor.4U cells (male) | Ncardia (previously Axiogenesis) | An internal line with no catalog number. Lot# MCB202CL |
| **Experimental Models: Organisms/Strains** | | |
| **Oligonucleotides** | | |
| **Recombinant DNA** | | |
| **Software and Algorithms** | | |
| Columbus Image Data Storage and Analysis System | PerkinElmer | http://www.perkinelmer.com/product/image-data-storage-and-analysis-system-columbus |
| edgeR | Robinson et al., 2009 | https://bioconductor.org/packages/release/bioc/html/edgeR.html |
| Gen5 | Biotek | https://www.biotek.com/products/software-robotics-software/gen5-microplate-reader-and-imager-software/ |
| G-means | Hamerly and Elkan, 2004 | N/A |
| goseq | Young et al., 2010 | https://bioconductor.org/packages/release/bioc/html/goseq.html |

| | | |
|---|---|---|
| ImageJ | NIH | https://imagej.nih.gov/ij/ |
| KinHub | Eid et al., 2017 | http://kinhub.org/ |
| Mathematica | Wolfram | https://www.wolfram.com/mathematica/ |
| Matlab | Mathworks | https://www.mathworks.com/products/matlab.html |
| Rstudio | The Rstudio team | https://www.rstudio.com/ |
| Seahorse Wave 2.4 Software | Agilent | https://www.agilent.com/en/products/cell-analysis/seahorse-wave-software |
| STAR aligner (via bcbio-nextgen) | Dobin et al., 2013 | https://github.com/bcbio/bcbio-nextgen |
| **Other** | | |
| Harvard Medical School Library of Integrated Network-based Cellular Signatures (LINCS) center dataset – RNAseq data | This paper | http://lincs.hms.harvard.edu/db/datasets/20324/ |
| Harvard Medical School LINCS center dataset – qRT-PCR data | This paper | http://lincs.hms.harvard.edu/db/datasets/20325/ |
